# N-glycosylation engineering in chimeric antigen receptor T cells enhances anti-tumor activity

**DOI:** 10.1101/2023.01.23.525164

**Authors:** Elien De Bousser, Nele Festjens, Evelyn Plets, Leander Meuris, Laura Vuylsteke, Daria Fijalkowska, Annelies Van Hecke, Elise Wyseure, Marie Goossens, Laure De Pryck, Sam Van den Bosch, Stijn De Munter, Bart Vandekerckhove, Nico Callewaert

## Abstract

Chimeric antigen receptor (CAR) T cell therapy has had limited success in solid tumors, requiring novel enhancement strategies. Modifying the glycocalyx of CAR T cells is unexplored; we report on genome-editing of the *MGAT5* gene to abolish human CAR T N-glycan poly-LacNAc modifications. This boosted tumor control in carcinoma and lymphoma models, for donors of whom the non-engineered CAR T cells largely failed in tumor control. More blood-circulating MGAT5 KO CD70 nanoCAR T cells were found, exhibiting potent tumor cell-killing activity *ex vivo*, while non-glycoengineered CAR T cells faltered. MGAT5 KO CD70 nanoCAR T cells also mediated durable anti-tumor immunity, improving control of secondary carcinoma challenge months later. Single-cell transcriptomics revealed increased mitotic activity and type I interferon signaling, indicating sustained intratumoral activation. The glyco-engineered cells had unaltered antigen sensitivity and dependence on T cell growth factors, preserving key safety features. *MGAT5* KO is readily compatible with clinical manufacturing, representing a promising approach to enhance CAR T cell therapy.

## Introduction

Immunotherapy with T cells that are genetically modified to express chimeric antigen receptors (CARs) has shown impressive efficacy in many, but by no means all patients suffering from a range of hematological malignancies^1–3^. The translation of this clinical success to a larger proportion of patients, and especially to the treatment of solid tumors, requires to overcome multiple obstacles^4,5^. In general, robust and stable populations of CAR T cells are required that are able to infiltrate the tumor and escape the immunosuppressive effect of the tumor micro-environment (TME), and which install long-term functional anti-tumor memory as to control also tumor relapses.

Whereas many aspects of T cell biology are being engineered to explore efficacy enhancement, the complex glycocalyx at the CAR T cell surface that mediates initial contacts of cells with their environment remains unexploited. However, basic research has revealed that surface glycosylation regulates important aspects of T cell functionality. For example, the expression of immune checkpoint inhibitors such as PD-1 and CTLA-4 is tuned by glycosylation^6,7^. Selective removal of NXS/T N-glycosylation sites from the TCR also enhances receptor diffusion, multimerization and functional avidity^8^. Furthermore, there is *in vitro* evidence that β-galactoside binding lectins (incl. mammalian galectins) can have a strong impact on the functionality of T cells. *Ex vivo* treatment of tumor-infiltrating T cells with an anti-galectin-3 antibody or a galectin competitive binder such as N-acetyllactosamine (LacNAc) resulted in the detachment of surface galectin-3, leading to increased cytotoxicity and ability to secrete cytokines such as IFN-γ^13,19^. Similarly, galectin-9 binding to TIM-3 was associated with the promotion of CD8^+^ T cell apoptosis^12,13^. On the other hand, galectin-1 was observed to promote expansion of regulatory T cells (Tregs) and induces apoptosis of specific thymocyte subsets and activated T cells, a process mediated at least in part through interaction with glycans on CD45^14,15^.

Inactivation experiments of the enzymes involved in the synthesis of poly-LacNAc structures corroborated their importance in T cell regulation. β1,6-N-acetylglucosaminyltransferase-V (MGAT5) is the enzyme responsible for the initiation of the GlcNAc-β-(1,6)-branch on N-glycans, which can be further elongated to carry poly-LacNAc chains (Figure 1.A). Absence of MGAT5, and thus a decrease in cell surface LacNAc density, lowers murine T cell activation thresholds by enhancing TCR clustering due to reduced galectin-glycoprotein lattice formation, which could be phenocopied *in vitro* by lactose addition to T cells^16^. Similarly, murine T cells mutated in β1,3-*N*-acetylglucosaminyltransferase (β3GnT2), which extends β1,6-branched *N*-glycans with poly-*N*-acetyllactosamine, show a lower threshold for TCR activation^17^ and proliferated more strongly than their wild type counterparts. More recently, the *MGAT5* gene was found as a key functionally validated hit in a CRISPR screen for enhanced intratumoral CD8^+^ T cell numbers in a murine model of glioblastoma^18^. With cessation of T cell responses, surface levels of the adhesion receptor cytotoxic T-lymphocyte antigen (CTLA-4) increase, a process dependent on increases in both CTLA-4 gene expression and MGAT5-mediated N-glycan branching^7^. Higher surface levels of CTLA-4 oppose CD28-dependent adhesion to CD86/CD80 on antigen-presenting cells, functioning as an immune checkpoint^19^. Furthermore, negative regulation of TCR signaling by β1,6-GlcNAc-containing N-glycans promotes development of Th2 over Th1 responses, enhancing Th2 polarization^20^. Moreover, *MGAT5* expression can be induced by the cytokine IL-10, decreasing antigen sensitivity of CD8^+^ T cells^21^. Many of these processes are involved in the immunosuppressive micro-environment of tumors. Accordingly, systemic inhibition in mice of the biosynthesis of LacNAc glycans via administration of a cell-membrane permeable enzyme inhibitor increased the number of infiltrating tumor-specific T cells^22^, as did administration of a polysaccharide with galectin-displacing properties^22,23^.

**Figure 1.**
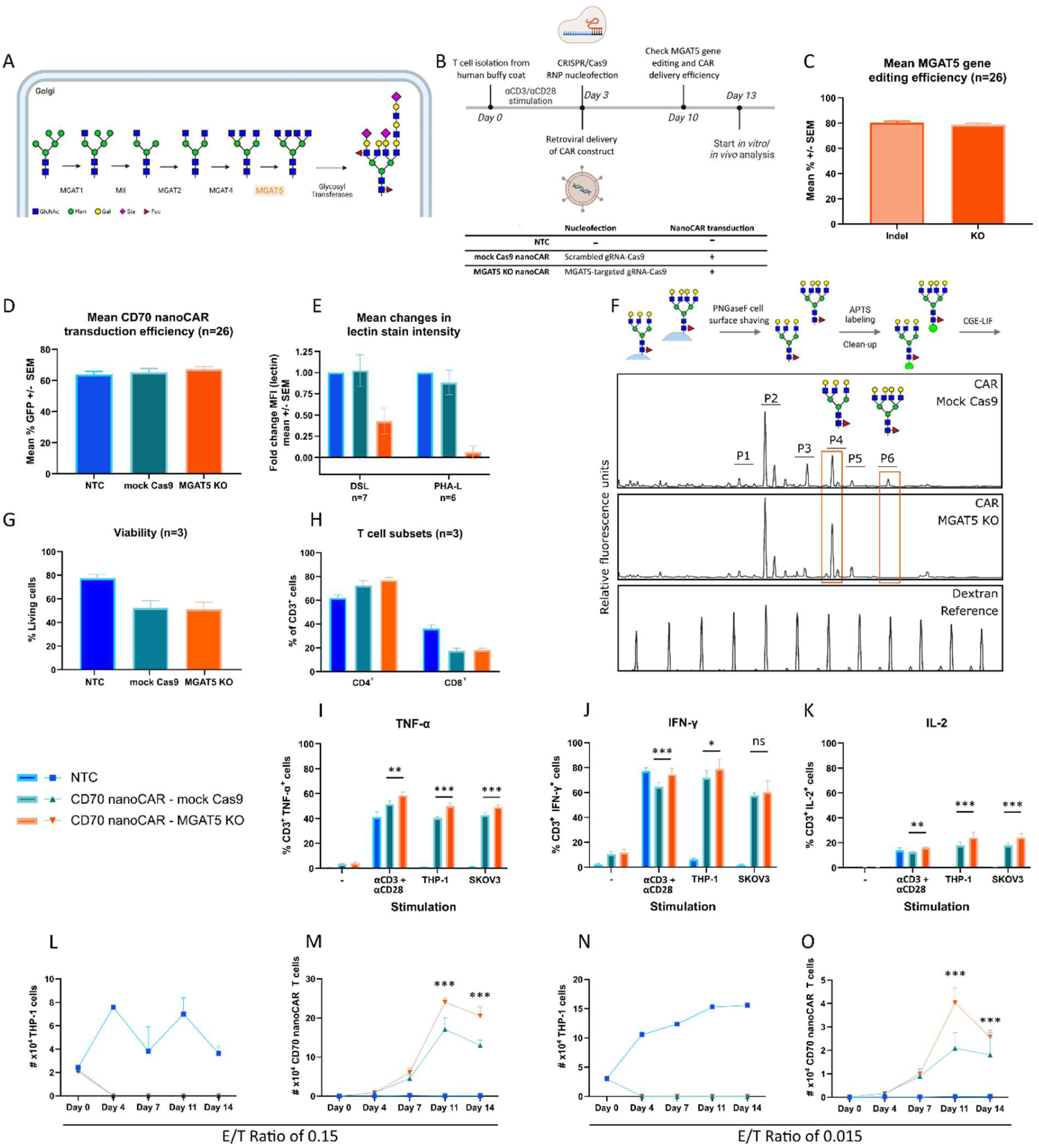
*MGAT5* glyco-gene editing and CD70 nanoCAR engineering and the impact of glyco-engineering in *MGAT5* on *in vitro* CD70 nanoCAR efficacy. A. Pathway of N-glycan branching. N-glycan branching is achieved through a series of mannosidase (M) and mannosylglycoprotein-N-acetylglucosaminyltransferase (MGAT)-mediated reactions. MGAT5, the enzyme of interest to our project is marked in orange. This branching primarily occurs in the medial-Golgi compartment. In later compartments, branched glycans are further modified by other enzymes including Gal-, GlcNAc-, sialyl- and fucosyltransferases to result in complex glycans. B. Experimental timeline. C. Mean editing efficiency as % insertions/deletions (indels) or proportion of indels that result in a frameshift or are 21+ bp in length (assumes all edits are in a coding region) (KO) obtained for the *MGAT5* locus. n=26, SEM: standard error of the mean. D. Mean transduction efficiency as % GFP expressing cells for the different engineering conditions. n=26, E and F. Profiling of alterations in cell surface glycosylation upon *MGAT5* KO in CD70 nanoCAR T cells. E. Lectin staining. 2 x 10^5^ wild type, mock-engineered (green) or *MGAT5* KO (orange) CAR T cells were collected, stained with fixable viability dye eFl780 and biotinylated lectin from *Datura stramonium* (DSL) or phytohemagglutinin-L lectin (PHA-L) followed by secondary staining with PE-coupled neutravidin. Analysis was done by flow cytometry and graphs show the mean reduction in lectin binding signal intensities as compared to wild type CAR T cells after gating on viable cells. Results are shown for engineered T cells from 7 (DSL)/6 (PHA-L) independent blood donors. F. CGE-LIF profiling. Schematic representation of the sample preparation and CGE-LIF profiles of the cell-surface glycome of mock-engineered and MGAT5 KO CD70 nanoCAR T cells. Sialidase digest was performed on the samples prior to the analysis. The major pairs of peaks in the profile are annotated P1-P6, based on the profile of total human plasma glycoprotein N-glycans which was run as a reference. The N-glycan profile of MGAT5 KO CD70 nanoCAR T cells is clearly different from that of mock-engineered CAR T cells. The peaks in P6 disappear while the peaks in P4 show a higher intensity relative to P2 and P3. This shift in electrophoretic mobility is consistent with the removal of one LacNAc unit (two monosaccharide units) or a shift from a tetra-antennary to a tri-antennary N-glycan. G. and H. Effect of *MGAT5* KO on CD70 nanoCAR viability and subset distribution at day 13 of *in vitro* culturing. At day 13, the immunophenotype of the cells was evaluated prior to the initiation of the functional assays. Mean data is shown from experiments performed with three independent donors. Error bars represent the SEM. I-K. The impact of glyco-engineering of *MGAT5* on *in vitro* CD70 nanoCAR cytokine production. Cytokine production of glyco-engineered CD70 nanoCAR T cells was evaluated by intracellular staining after co-incubation with THP-1 and SKOV-3 target cell lines for 16 hours. Unstimulated cells were included as negative control (-) while Immunocult stimulation was included as positive control (+). Technical duplicates were analyzed. The bars show the mean fractions of TNF-α, IFN-γ and IL-2 positive CD3^+^ obtained from three independent donors. Error bars represent the standard deviation. L-O. The impact of glyco-engineering in *MGAT5* on *in vitro* CD70 nanoCAR cytotoxic potential. Glyco-engineered CD70 nanoCAR T cells cultured in the presence of IL-7 and IL-15 were incubated at different effector to target THP-1 cell ratios (E/T) in technical duplicates and cell numbers were analyzed over a time period of 14 days. A second challenge with THP-1 cells was added at day 7. Error bars represent the SEM on cell number from data obtained with 3 different T cell donors. L and M. Results for E/T ratio of 0.15 corresponding to the co-culture of 20 000 THP-1 cells with 3 000 glyco-engineered CD70 nanoCAR T cells. N and O. Results for E/T ratio of 0.015 corresponding to the co-culture of 20 000 THP-1 cells with 300 CD70 nanoCAR T cells. To account for the correlation over time, the data from panels M and O was modeled using a mixed negative binomial model with a random intercept for each cluster (a cluster being defined as the set of measurements sharing the same donor, E/T Ratio and type of CAR T cells). We found that, on day 11, the number of *MGAT5* KO nanoCAR T cells is about 1.74 times higher (95% CI: 1.36 to 2.21) than the number of nanoCAR T cells starting from the same conditions. On day 14, the number of *MGAT5* KO nanoCAR T similarly is about 1.70 times higher (95% CI: 1.33 to 2.18). All these estimates are averaged over the two E/T ratios and the three independent donors; *** p < 0.001, ** 0.001 < p < 0.01, * 0.01 < p < 0.05, ns p >0.05

Taken together, these fundamental immune-glycobiological study data gathered over two decades from multiple labs on poly-LacNAc modifications on T cells, indicated that poly-LacNAc modification in the T cell glycocalyx may function as a multipronged checkpoint mechanism that limits antitumor T cell immunity. Glycobiology can be notoriously difficult to translate from murine studies to humans, as qualitative and quantitative aspects of the outer branch modifications of the glycocalyx are amongst the most rapidly evolving mammalian cellular structures, likely under differential pathogen pressure^24^. Whether sufficient aspects of the mainly murine T cell based glycobiology described above would be applicable to human CAR T cells, with their independence from TCR-antigen recognition and the intricacies of TCR-coreceptor clustering, was unexplored. Hence, we set out to test the hypothesis whether genetic inactivation of poly-LacNAc N-glycan biosynthesis in human CAR T cells by *MGAT5* gene inactivation would lead to more efficacious treatment of human lymphoma and carcinoma tumor models. These studies are the first to report on targeting of the glycocalyx of CAR T cells for therapy enhancement.

## Results

### Generating MGAT5 knockout (KO) CD70 nanoCAR T cells

Staying close to the currently used methods for human CAR T cell therapy manufacturing, we optimized a workflow for the combined CRISPR-Cas9 mediated *MGAT5* mutagenesis and retroviral CAR delivery to purified, activated human CD3^+^ T cells (Figure 1.B and Supplementary Figure 2). The presence of both CD4^+^ and CD8^+^ T cell subsets in the final CAR T cell product is indispensable for efficient anti-tumor immunity^25^. The mean editing efficiency as percentage insertions and deletions (% indel) for the *MGAT5* locus over multiple (n=26) experiments was consistently high (exceeding 80% indel) as is depicted in Figure 1.C. Flow cytometry was used to measure both CD70 nanoCAR expression and GFP expression as readouts of the retroviral transduction efficiency^26^. CD70 nanoCAR transduction efficiencies of more than 70% were consistently obtained over multiple (n=26) experiments, irrespective of whether the *MGAT5* gene was engineered or not (Figure 1.D).

### MGAT5 KO CD70 nanoCAR T cells have strongly reduced N-glycan poly-LacNAc cell surface density

In order to assess the extent of the intended reduction in poly-LacNAc cell surface abundance, we used flow cytometry to measure binding of the lectin from *Datura stramonium* (DSL) (Figure 1.E). This lectin binds to all glycans containing (poly-)LacNAc glycotopes regardless of whether they are linked to N-glycans, O-glycans or glycosphingolipids. MGAT5 KO CD70 nanoCAR T cells had about 50% reduced staining with DSL. Furthermore, using the lectin from *Phaseolus vulgaris* (PHA-L), which specifically binds to N-glycans carrying the MGAT5-synthesized β-1,6 branch, we observed near-complete loss of binding (Figure 1.E). Together, these results show that the extent of *MGAT5* genome editing efficiency that we achieved successfully eliminates N-glycan β1,6-branching. The loss of β-1,6 branch formation on the N-glycans was corroborated by direct structural profiling of the N-glycans shaved off the cell surface using PNGase F enzyme, followed by capillary gel electrophoresis with laser induced fluorescence (CGE-LIF) (Figure 1.F), a method previously developed at our lab^27,28^. The desialylated N-glycan profile of MGAT5 KO CD70 nanoCAR T cells demonstrates a loss of the main tetra-antennary N-glycan species (synthesis of which requires MGAT5), whereas the tri-antennary precursor N-glycans that MGAT5 takes as its substrates are increased in relative abundance, as expected.

### MGAT5 KO CD70 nanoCAR T cells are fully functional *in vitro* and expand more upon repeated tumor cell challenge

MGAT5 KO CD70 nanoCAR T cells have equivalent viability rates as non-glycoengineered control cells *in vitro* (Figure 1.G). After CAR construct transduction and subsequent cultivation, most of the CAR T cells in the total CD3^+^ T cell pool are CD4^+^, and this was even more pronounced when CD70 nanoCAR T cells underwent CRISPR-Cas9 engineering (both for mock gRNA and *MGAT5* KO gRNA), generating a CAR T cell preparation with about 80% CD4^+^ and 20% CD8^+^ (Figure 1.H). Using flow cytometrical methods, we quantified production proficiency for the TNF-α, IFN-γ and IL-2 cytokines of the glyco-engineered CD70 nanoCAR T cells, after challenging them with the THP-1 and SKOV-3 target tumor cell lines (Figure 1.I-K). At least as high a proportion of the MGAT5 KO CD70 nanoCAR T cells are able to produce cytokines upon antigen stimulation as are mock-nucleofected CD70 nanoCAR T cells. This cytokine expression is dependent on CD70 nanoCAR expression, given that the fraction of cytokine-expressing non-transduced T cells (NTC) is close to zero or very low in the presence of CD70 positive THP-1 cells (but high after polyclonal Immunocult stimulation). In order to evaluate the combined proliferative and cytotoxic potential of MGAT5 KO CD70 nanoCAR T cells, very low proportions of CAR T cells were co-cultured with THP-1 target cells. The number of THP-1 cells left surviving in culture was determined by flow cytometry. At day 7, a second challenge with target THP-1 cells was performed. Figure 1.L and N show the results corresponding to an effector/target (E/T) ratio of 0.15 and 0.015. At these ratios, all target cells get killed by day 4, in all of the mock-nucleofected and MGAT5 KO CD70 nanoCAR T cell conditions. Interestingly, upon second challenge with tumor cells, at both E/T ratios, the number of CD70 nanoCAR T cells is significantly higher for those that are KO in *MGAT5*, indicating a stronger and more sustained antigen recognition-induced expansion of these glyco-engineered CAR T cells (Figure 1.M and O).

Treatment of carcinoma-bearing mice with MGAT5 KO CD70 nanoCAR T cells yields improved tumor control A key challenge with translationally relevant CAR T cell studies using human donor cells in preclinical mouse models is the substantial observed biological variability from donor to donor in the derived baseline CAR T cell potency^29–31^, varying from complete tumor control to barely any tumor control, which is also observed in the clinic. Hence, also our studies aimed at realistically preclinically evaluating human CAR T MGAT5 engineering in human tumor xenograft mouse models. This required a significant experimental effort to obtain a representative picture of the spectrum of effect sizes of the *MGAT5* KO engineering, observed over 3 or 4 donors each time.

As CAR T cell therapy clinically so far mostly fails in carcinomas and improvements there would open exciting translational opportunities, we chose to first test experimental therapeutic efficacy of the MGAT5 KO CD70 nanoCAR T cells in the NSG^32^ mouse xenografted with CD70-expressing SKOV-3 ovarium carcinoma cells, the latter engineered to produce luciferase, for longitudinal tumor burden measurement. These tumors establish a micro-environment in which most of the known T cell immunosuppressive mechanisms are at play, thus forming a truly challenging target for CAR T therapy^33–36^. The experimental layout is illustrated in Figure 2.A and runs for 120 days, including primary tumor challenge and relapse-mimicking secondary tumor challenge in surviving mice, to assess long-term functional persistence of CAR T cells. Full time-course results were obtained for three independent experiments, performed with T cells from different donors (Donor A, B and C). The outcome of treatment for the primary tumor was categorized as: (1) full control, i.e. the tumor becomes undetectable and no relapse follows; (2) full control but occurrence of a relapse later on; (3) partial control, i.e. a halt in tumor growth but the tumor remains detectable; (4) no control of tumor growth throughout the duration of the experiment.

**Figure 2.**
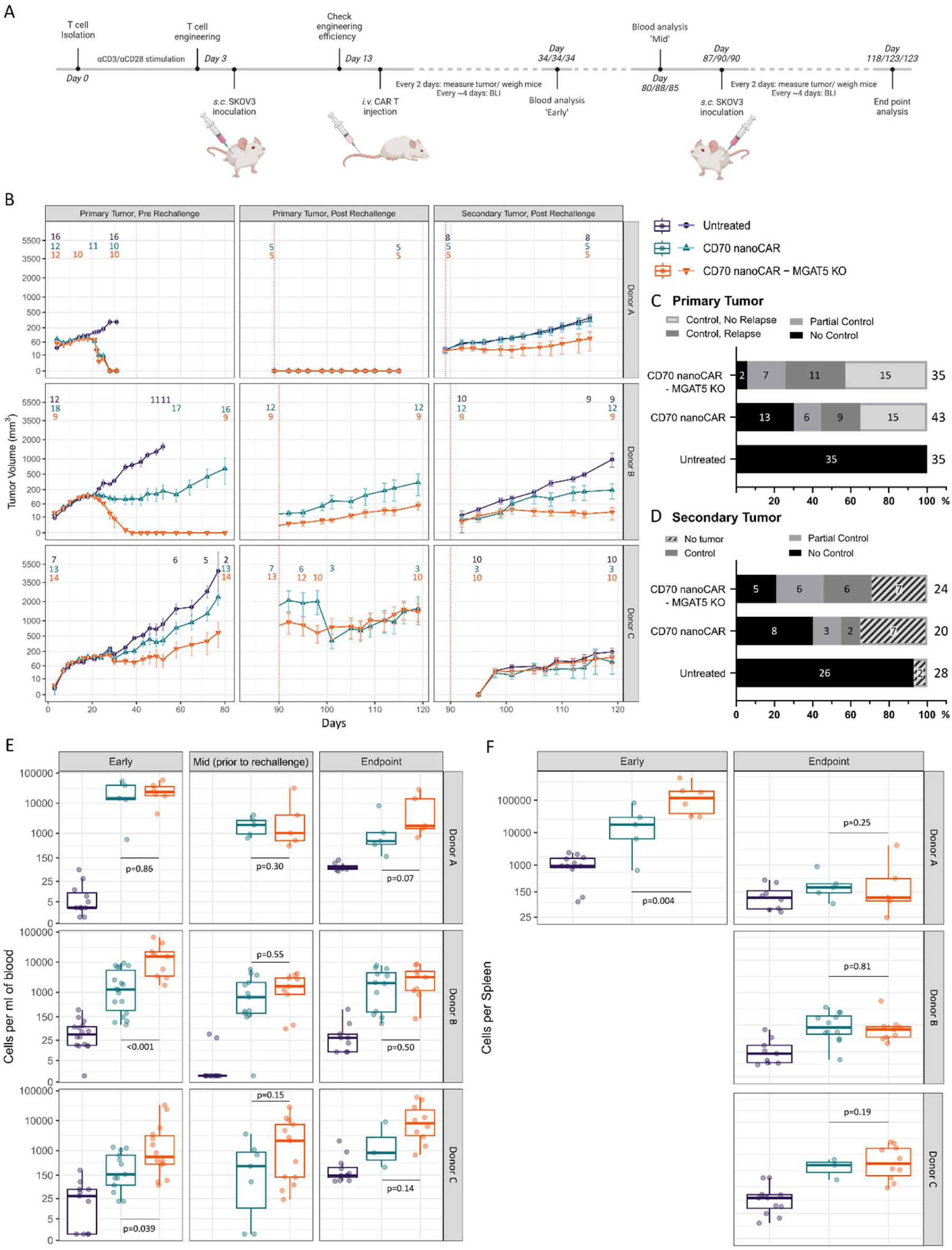
The impact of glyco-engineering on *in vivo* CD70 nanoCAR functionality in the SKOV-3 solid tumor model. A. Schematic representation of the experimental timeline for the study of the *in vivo* efficacy of MGAT5 KO CD70 nanoCAR T cells. B. Mean tumor burden measured by caliper. Tumor volume is calculated as (tumor length x tumor width^2^)/2. Group means are indicated with error bars representing the SEM. Original values of tumor volume are shown, Y-axis is cube root transformed. Number of mice surviving in each group are indicated above the datapoints and adapted at timepoints whenever a mouse was sacrificed at humane endpoint. Timepoint of rechallenge is indicated with at dashed red line. C. Overview of the response to primary tumor challenge in the different treatment groups for all three experiments combined. D. Overview of the response to secondary tumor challenge in the different treatment groups for all three experiments combined. E and F. Number of CAR T cells identified in the different treatment conditions in peripheral blood (E) and spleen (F) at day 34/34/34 (Early), day 80/88/85 (Mid) and day 118/123/123 (Endpoint) in three independent experiments with cells derived from donors A/B/C. Data is represented as concentration of CD3^+^GFP^+^ cells per ml of blood or per spleen. The number of CAR T cells in the spleen is counted based on a gating strategy that includes both CD4^+^ and CD8^+^ T cells. Each data point represents a single animal. The difference between groups was tested using a quasi-Poisson GLM at each timepoint and for each donor, p-values are indicated within the respective plots.

As is clear from Figure 2.B and the bar graphs in Figure 2.C, in this model the primary tumor is never controlled by the mice that did not receive CAR T cells; they were all sacrificed at the humane endpoint. A considerable number of mice (n=19) in the non-glycoengineered CAR T cell treated groups across the 3 donors show only partial or even no control at all of the primary tumor, while this was the case for only 9 mice in the MGAT5 KO groups.

Donor A represents the case in which potent CAR T cells can be obtained without any further engineering: even the non-glycoengineered CAR T cells were fully efficacious at controlling the primary tumor and we did not observe any relapse at the primary tumor site over time (Figure 2.B and Supplementary Figure 4.A-B). Donors B and C represent the clinically more frequently observed picture where CAR T cells have variable and generally low capacity of clearing carcinoma. For both donors, MGAT5 KO CD70 nanoCAR T treatment led to significantly improved primary tumor control compared to treatment with non-engineered CD70 nanoCAR T cells (Supplementary Figure 5.A-B and Supplementary Figure 6.A-B). However, for these donors the primary tumor regrew even when it was initially cleared (Figure 2.B). Using the data for these two donors where relapses were observed, a Kaplan-Meier survival analysis detected that a longer relapse-free survival was obtained in the MGAT5 KO CD70 nanoCAR T treated groups (Supplementary Figure 3.A-B).

To test for long-term functional persistence of the CAR T cells and capability of mounting a functional anti-tumor response, we mimicked a relapse tumor after a period of approximately 90 days, by rechallenge with a secondary tumor in the opposite (left) flank of the CAR T treated mice that survived the primary tumor. For this secondary tumor, we defined four types of tumor control: (1) no tumor, meaning the secondary tumor never develops; (2) full control, i.e. the secondary tumor becomes undetectable after an initial growth phase; (3) partial control, i.e. the secondary tumor stops growing but remains detectable; (4) no control of secondary tumor growth throughout the duration of the experiment. In order to have a control group (untreated) with comparable age and history as the remaining mice from the treatment arms, we tumor-challenged a group of mice that were kept in husbandry from the very start of the experiment but that did not receive a primary tumor challenge. As shown in Figure 2.B and the bar graph in Figure 2.D, MGAT5 KO CD70 nanoCAR T cell treatment also leads to better rechallenge tumor control. In the experiment with CAR T cells from donor A, the average tumor volume at the end of the analysis is an estimated 6.4 times smaller (p=0.011) in mice that were treated with MGAT5 KO CD70 nanoCAR T cells compared to non-glycoengineered CAR T treated mice (Supplementary Figure 4.C-D). For those mice treated with CAR T cells from donor B that still had a remnant of the primary tumor at the timepoint of rechallenge, we found that at the end of the experiment, these tumors had a smaller average size in the MGAT5 KO group than in the non-glycoengineered CAR T cell treated group, although the difference was not statistically significant (Figure 2.B, Supplementary Figure 5.C-D).

Hence, we concluded that in all analyses where non-engineered donor CAR T cells did not or partially control the tumors, their MGAT5 KO-engineered counterparts substantially rescued this.

When MGAT5 KO enhances primary tumor control, higher numbers of MGAT5 KO CD70 nanoCAR T cells are present in blood as compared to non-glycoengineered CD70 nanoCAR T cells

We quantified the number of circulating human CD3^+^ T cells and the CAR-engineered subpopulation thereof in blood. We first sampled blood on day 34, which is 21 days after CAR T cell administration. At this timepoint, the spectrum of donor-dependent variance in therapeutic response is first clearly detected, (Figure 2.B, left panels), making it a suitable timing to measure the strength of the CAR T proliferation.

Interestingly, for the two donors (B and C) whose non-glycoengineered CAR T cells did not completely control the tumor and where MGAT5 KO CD70 nanoCAR T cells yielded substantial tumor control enhancement, the number of MGAT5 KO CD70 nanoCAR T cells in blood (Figure 2.E, left panels) is significantly increased as compared to the counterpart non-glycoengineered CD70 nanoCAR T cells. In contrast, for donor A, whose CAR T cells completely eradicated the tumor already without glyco-engineering, even higher numbers of CAR T cells were present already for non-engineered CAR T cells and this number was not further boosted by *MGAT5* KO engineering. It is also striking that in donor C, where CAR T therapy was the least successful, also the lowest average number of CAR T cells amongst the 3 donors are found. *MGAT5* KO boosts this significantly leading to strongly enhanced survival of the challenged mice (at day 84 just prior to second tumor challenge: 13 out of 14 surviving for MGAT5 KO CD70 nanoCAR T vs. 7 out of 13 for non-glycoengineered CAR T), with 10 also surviving secondary tumor challenge, whereas only 3 mice were still alive in the non-glycoengineered CAR T treated group at the end of the experiment. Donor B is intermediate between A and C and indeed also had intermediate levels of circulating non-engineered CAR T cells. The *MGAT5* KO boost of this donor’s CAR T cells was here sufficient to get all primary tumors cleared initially, with full survival (9 out of 9 mice) till day 90 (when we performed tumor rechallenge), whereas by this timepoint we lost 1/3^rd^ of the mice treated with non-engineered CAR T cells (12 out of 18 surviving). Hence, this exploration revealed that there is a striking correspondence between the increased blood concentration of CAR T cells and the tumor control success.

Blood sampling was repeated around day 80 (i.e. shortly prior to tumor rechallenge) and at the end of the experiments on days 118-123 (which is about 35 days after the tumor rechallenge), at which time also the spleen CAR T cell counts of all mice surviving up to that point were measured. At the late timepoint prior to tumor rechallenge (day 80), a more than 10-fold contraction of the concentration of CAR T cells was observed for donor A (Figure 2.E, compare right and middle panels) where no tumor had been present for about 2 months, as one would expect in the absence of CAR antigen. In donor B, such contraction was only observed for the MGAT5 KO group and to a lesser extent than in donor A; this is in line with the observation that this group had cleared the tumor initially, but very small tumors slowly regrew, providing some antigenic stimulation. In contrast, the CAR T numbers in the group treated with non-engineered donor B CAR T cells did not contract, in accordance with most surviving mice never having cleared their tumor over this time course, providing continuous antigenic stimulation. The same was seen in donor C, here both for the non-glycoengineered CAR T group and for the MGAT5 KO CD70 nanoCAR T group, in line with continuous tumor exposure in the mice.

Upon tumor rechallenge (Figure 2.E, compare middle and right panels) no major differences occurred in donor B, whereas in donor A, MGAT5 KO CD70 nanoCAR T blood cell concentration appeared to somewhat re-expand by this renewed antigen stimulation vs. a further mild contraction observed for non-glyco-engineered cells, with the MGAT5 KO treated group indeed achieving better secondary tumor control (Figure 2.B, right panels). It must be noted that all of these median levels for donor A and donor B are still around or above 1000 CAR T cells per ml of blood, which also at the day 34 timepoint appears to be close to the threshold needed to achieve SKOV-3 tumor control. Remarkably, for donor C, rechallenge with the tumor also leads to further expansion of the MGAT5 KO CD70 nanoCAR T cells, to the levels seen to robustly eliminate tumor cells in the other donors. However, no differences could be observed in terms of tumor killing (Supplementary Figure 6.C-D).

For the experiment with donor A in which the primary tumor was completely cleared, also with non-glycoengineered CAR T cells, we sacrificed half of the mice at day 34 in order to investigate spleen CAR T counts at that early timepoint as well. Remarkably, the number of MGAT5 KO CD70 nanoCAR T cells in the spleen was higher in these mice vs. the non-glycoengineered CAR T cells (Figure 2.F), and the numbers were extremely high (median above 100,000 CAR T cells per spleen for the MGAT5 KO group). Investigating the spleens at the end of the experiments, there were no longer differences in CAR T cell numbers between MGAT5 KO and non-glycoengineered groups, for all donors, and the numbers were much lower, close to background for donor A, intermediate for donor B and highest for donor C (medians not exceeding 10,000 cells per spleen).

MGAT5 KO CD70 nanoCAR T cells enhance control of a highly aggressive metastatic B cell lymphoma model To investigate the scope of applicability of the MGAT5 KO engineering module, we have expanded our findings to a very different tumor type, i.e. the Non-Hodgkin B cell lymphoma setting in which CAR T therapy is already part of the standard of care treatment paradigm, but where efficacy enhancement is still also sorely needed. In contrast to the localized subcutaneous SKOV-3 tumor, in this case NSG mice were intravenously injected with luciferase-expressing Raji cells, resulting in a metastatic model that establishes a very rapidly growing multi-systemic human lymphoma burden, incl. in the brain^37^. This forms a stringent test for CAR T cell therapy, as therapeutic cells need to exert tumor control very rapidly, which leads to terminal tumor burden in all mice within 1 week after the treatment timepoint when left untreated. This is translationally reminiscent of the clinical setting of treating patients with relapsing lymphoma. Raji lymphoma also expresses the CD70 antigen (Supplementary Figure 1), which is an emerging new target antigen in both B and T cell lymphomas also clinically^38,39^, so we could use the same nanoCAR against CD70 that we used for the above SKOV-3 carcinoma work, allowing for comparison of the results. Seven days after tumor cell inoculation, mice were treated with either mock Cas9-engineered or MGAT5 KO CD70 nanoCAR T cells. As control groups, mice were treated with non-transduced T cells to evaluate tumor development. Throughout the experiment, tumor burden was measured by bioluminescence imaging (BLI). A schematic representation of the experimental timeline is depicted in Figure 3.A. The experiment has been repeated on four different donors (donor D, E, F, G), in order to be able to take into consideration the variability in the efficacy of CAR T cells derived from different donors. For two out of four donors (F and G) we found that *MGAT5* KO leads to a >50% enhanced long-term survival rate (Figure 3.B), with monitoring in all experiments for at least 90 days post-tumor challenge. As with the SKOV-3 carcinoma model, again we observed that in these two donors where MGAT5 KO resulted in the biggest enhancement vs. mock-engineered CAR T cells, a significantly higher number of MGAT5 KO CD70 nanoCAR T cells were found in the blood at the early timepoint 2 weeks after CAR T treatment (Figure 3.C). In most of these long-term surviving mice, the tumor became undetectable (Figure 3.B-C, right panels), a truly remarkable result given the extreme aggressivity of this tumor model. For donor E, the non-engineered CAR T cells already yielded survival of 3 out of 4 mice, and MGAT5 KO turned this into complete protection (4/4). Here, we also found a trend to higher numbers of MGAT5 KO CD70 nanoCAR T cells in the blood. In donor D, neither non-engineered nor MGAT5 KO CD70 nanoCAR T cells yielded any long-term survival. This donor’s cells were an outlier, with massive CAR T cell proliferation, yielding cell numbers at least 10-fold higher than in any of our other experiments. It appears that these donor’s CAR T cells are extraordinarily stimulated to proliferate, yet defective in tumor killing capacity.

**Figure 3.**
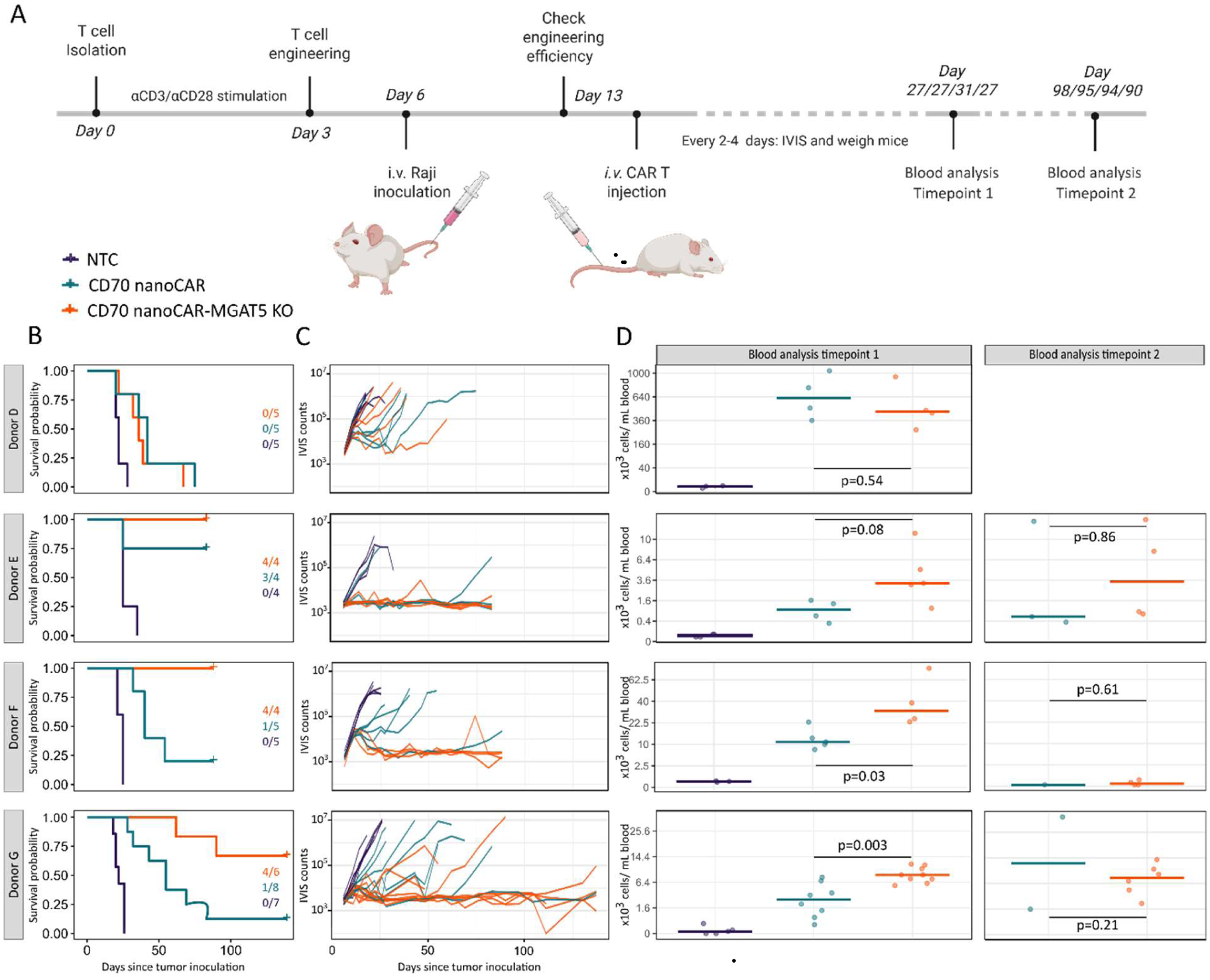
The impact of *MGAT 5* KO on *in vivo* CD70 nanoCAR functionality in the Raji lymphoma model. A. Schematic representation of the experimental timeline for the study of the *in vivo* efficacy of MGAT5 KO CD70 nanoCAR T cells. B Kaplan-Meier survival curves for CD70 nanoCAR treated mice. The number of surviving mice at the end of the experiment vs. the number of mice per group at the start of the experiment are indicated. Data from 4 independent donors (D, E, F, G) are shown. C. Bioluminescence imaging of tumor burden in all mice individually. IVIS counts are total whole-body counts measured as photons/sec. D. Flow cytometry-based analysis of CD70 nanoCAR T cells in blood. Number of CAR T cells identified in the different treatment conditions in peripheral blood at day 27/27/31/27 (Timepoint 1) and day 98/95/94/90 (Timepoint 2), in four independent experiments with cells derived from donors D/E/F/G. CAR T cells were detected as human CD3^+^GFP^+^ cells. Each data point represents a single animal. The number of CAR T cells present in the blood is indicated as cells/mL blood. The difference between groups was tested using a quasi-Poisson GLM at each timepoint and for each donor, p-values are indicated within the respective plots.

In conclusion, MGAT5 KO substantially enhanced the protective capacity of CD70 nanoCAR T cells in the metastatic Raji lymphoma model, concomitant with enhanced concentration of CAR T cells in the blood, therewith corroborating the results obtained in the SKOV-3 model as described above.

The higher number of circulating MGAT5 KO nanoCAR T cells results in higher *ex vivo* tumor cell killing activity over prolonged tumor cell challenge.

From these findings, the simplest conceivable mechanism of action of the MGAT5 KO-mediated CAR T cell enhancement was a higher and/or more prolonged circulating therapeutic cell level available to combat tumor growth, resulting from stronger proliferation and/or from stronger tumor cell killing potency of the CAR T cells. For this, it was necessary to investigate whether this enhanced concentration of circulating MGAT5 KO CD70 nanoCAR T cells in blood is indeed competent by itself at more effectively killing tumor cells *ex vivo* (Figure 4). Hereto, we prepared blood from SKOV-3 tumor-bearing mice 21 days after treatment with either non-glycoengineered CD70 nanoCAR T cells versus MGAT5 KO CD70 nanoCAR T cells from 2 extra independent donors (donor H/I) (Figure 4.A). For donor H, SKOV-3 tumor stasis was achieved in the mice in the MGAT5 KO CD70 nanoCAR T treated group, while this was not the case for the mock CD70 nanoCAR T treated group (Figure 4.B), similar to the case of donor C described above (Figure 2). For donor I, the MGAT5 KO and mock engineered CD70 nanoCART cells both completely cleared the tumor (Figure 4.F), similar to donor A above (Figure 2). In both cases, higher numbers of MGAT5 KO CD70 nanoCAR T cells were again detected (Figure 4.C and G). To investigate whether this resulted in higher tumor cell killing activity per volume of blood, cells prepared from equivalent volumes of blood were co-cultured with THP-1 tumor cells (which are CD70 expressing suspension cells and thus more amenable to this co-culture analysis than SKOV-3) at different ratios, and THP-1 cell counts as well as CAR T counts were followed over time. Figure 4.D shows that circulating MGAT5 KO CD70 nanoCAR T cells from donor H efficiently and almost completely wiped out the THP-1 tumor cells, even upon a second THP-1 tumor cell challenge at day 7, whereas blood cells from the non-glyco-engineered CAR T treated mice were ineffective at controlling the THP-1 cells even in the first round of challenge. Circulating non-glyco-engineered CAR T cells from donor I showed delayed control of THP-1 cells compared to MGAT5 KO CD70 nanoCAR T cells (Figure 4.H), with viable THP-1 cells still observed at day 4 of co-culture, whereas at day 7 both blood-derived CAR T preparations had cleared the tumor cells. There was no control of THP-1 target cells when using blood cells from mice treated with non-engineered T cells from the same donors. Moreover, MGAT5 KO CD70 nanoCAR T cells from both donors proliferated much stronger than non-engineered counterparts in these *ex vivo* co-cultures, and in both cases sustained this also after the second THP-1 cell addition (Figure 4.E and I). Hence, we conclude that MGAT5 KO results in a higher level of tumor cell killing-competent CAR T cells surveilling the body of the CAR T treated mice, with these cells remaining cytotoxic and proliferating also after prolonged tumor cell encounter, providing a simple mechanism for the enhanced observed *in vivo* tumor control.

**Figure 4.**
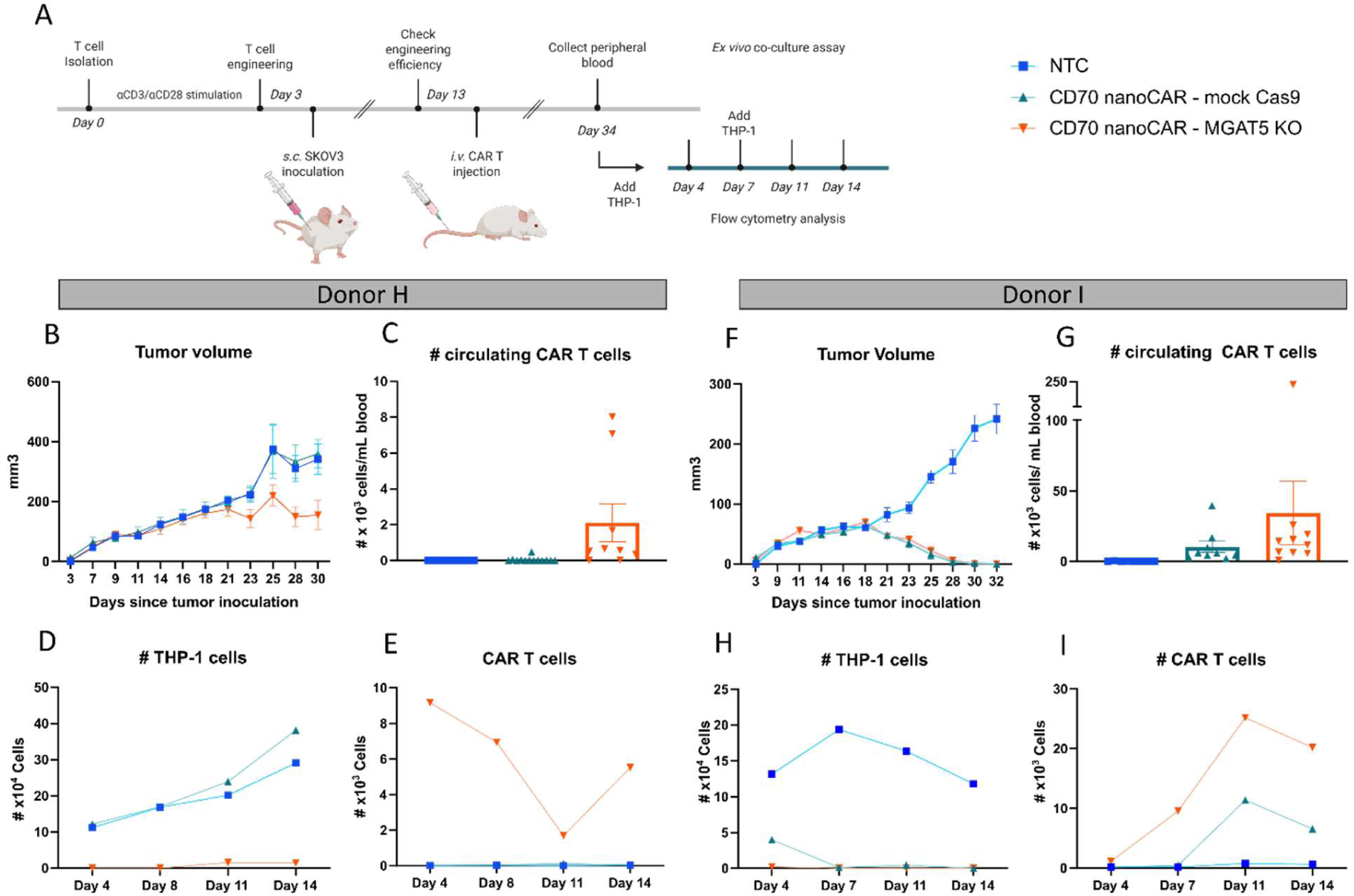
The impact of glyco-engineering in *MGAT5* on *ex vivo* CD70 nanoCAR efficacy. A. Time line of the experiment. Data obtained for cells engineered from donor H (panel B-E) and donor I (panel F-I). B and F. SKOV-3 tumor burden measured by caliper. Tumor volume is calculated as (tumor length x tumor width2)/2. Group means are indicated with error bars representing the SEM. C and G. The number of CD70 nanoCAR T cells (represented as number of CD3^+^GFP^+^ cells) present in the blood is indicated as cells/mL blood. Each data point represents a single animal. Error bars represent the SEM. D-E and H-I. The impact of glyco-engineering in *MGAT5* on *ex vivo* CD70 nanoCAR cytotoxic potential. Blood was prepared from SKOV-3 tumor bearing mice 21 days after treatment with either WT or MGAT5 KO CD70 nanoCAR T cells. Cells prepared from 50µL blood (donor H) or 40µL blood (donor I) were put in co-culture with 20 000 THP-1 cells and THP-1 and CAR T cell counts were followed over a time period of 14 days. A second challenge with THP-1 cells was added at day 7.

MGAT5 KO CD70 nanoCAR T cells require equal antigen density to degranulate and remain fully dependent on cytokine growth factors for their proliferation.

Beyond elevated concentrations of circulating CAR T cells with persistent tumor cell killing competence, another conceivable mechanism of action for MGAT5 KO in enhancing antitumoral potency could be a lowered threshold of antigen density required for CAR T cell activation. To investigate this, we have analyzed degranulation levels (using the CD107a/LAMP1 exposure on the cell membrane as a proxy) in function of incubation of CAR T cells on varying concentrations of coated recombinant human CD70 antigen. We find similar activation thresholds for non-glycoengineered versus MGAT5 KO CD70 nanoCAR T cells, indicating that increased tumor cell control of MGAT5 KO CD70 nanoCAR T cells is not due to a decreased activation threshold, which otherwise may increase the risk for overstimulation or off-target activation (Figure 5.A).

**Figure 5.**
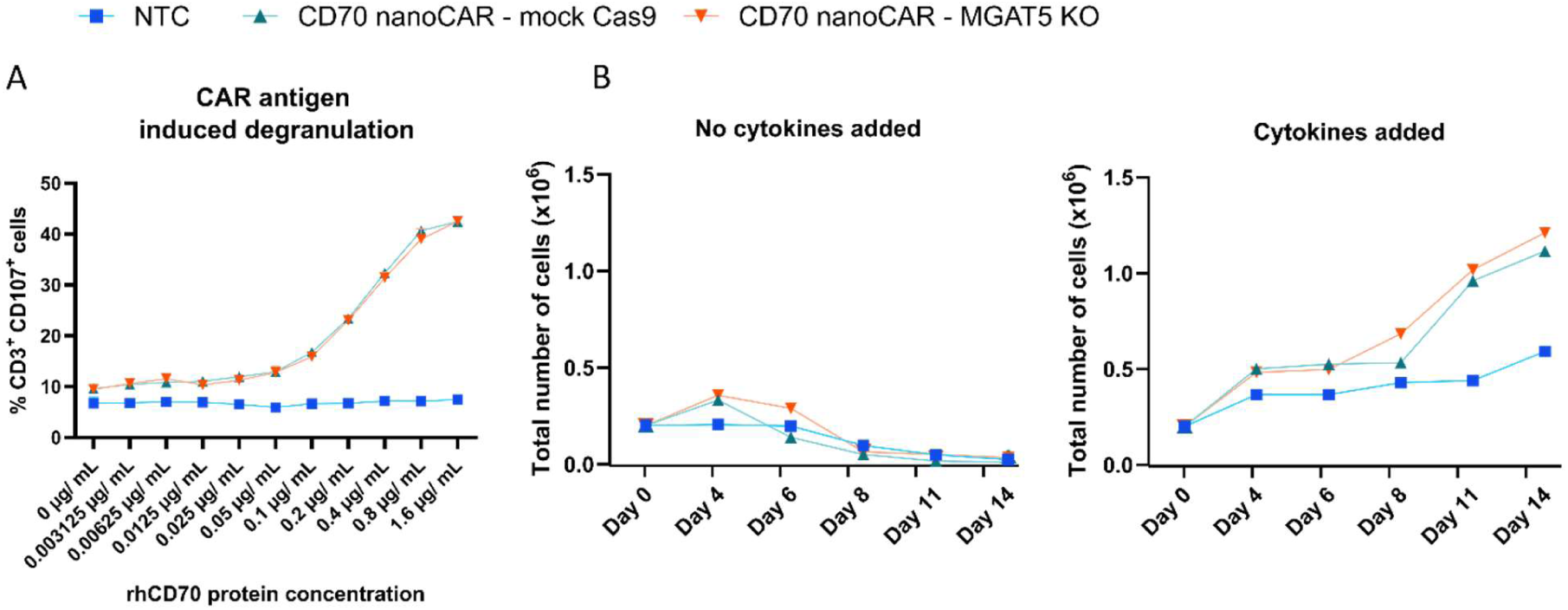
MGAT5 KO CD70 nanoCAR T cells require equal antigen density to degranulate and fully depend on cytokine growth factors for their proliferation. A. Flow cytometry analysis of CD107a degranulation assay, on cells from donor K. Untransduced T cells (NTC) and CD70 nanoCAR T cells with and without *MGAT5* engineering were co-cultured in technical duplicate with different concentrations (3.125 ng/ mL up to 1.6 µg/ mL) of coated rhCD70 protein for 5 hours. CD107a^+^ cell staining was analyzed by flow cytometry gating on CD3^+^ cells. Data shown is representative of three experiments performed with T cells obtained from two independent donors. B. Cytokine independent growth assay performed on cells engineered and cultured in the presence of IL-7 and IL-15 as described in the engineering protocol (from donor G). Left, total T cell count of engineered T cells cultured in the presence IL-7 and IL-15 for an additional 14 days. Right, total T cell count of engineered T cells cultured in the absence of supplemental cytokines.

Furthermore, in the absence of antigen, neither mock nor MGAT5 KO CD70 nanoCAR T cells grow without IL-7/IL-15 (growth/survival-associated cytokines) (Figure 5.B), demonstrating that these nanoCAR T cells are still signal-3 dependent and not transformed, a key safety requirement. Together, the data indicate that MGAT5 KO CD70 nanoCAR T cells retain an inherent capacity for more vigorous and prolonged antigen- and cytokine-dependent proliferation without making the cells hypersensitive to antigen.

Furthermore, we did not observe any signs of graft-versus-host-disease (GvHD) in any of the SKOV-3 *in vivo* experiments with 5 human donors. In the Raji model, in which CAR T cells proliferate to much higher numbers than in the SKOV-3 model, we observed signs of GvHD in 1 out of 22 mice in the non-glycoengineered CAR T treatment groups across the 4 donor experiments, and in 3 out of 19 mice in the MGAT5 KO nanoCAR T treatment groups.

MGAT5 KO enhances type I IFN signaling and other hallmarks of enhanced persistent activation and proliferation in tumor-infiltrating CAR T cells What remained was to explore which alterations were present in the state of MGAT5 KO tumor-infiltrating nanoCAR T cells. To this effect, 10 days after SKOV-3 tumor inoculation, we treated groups of 6 mice with CD70 nanoCAR T cells and after 7 days we collected peripheral blood and resected the tumors. CAR T cells were sorted from these samples and we performed single-cell RNA sequencing (scRNAseq) (Figure 6.A). As can be seen in Figure 6.B, this timepoint 17 days post-tumor inoculation is a few days before we start measuring tumor size divergence between non-glycoengineered and MGAT5 KO CD70 nanoCAR T cell treated mice, and was chosen as it should allow for the most unbiased sampling of initial tumor attack. Uniform manifold approximation and projection (UMAP) analysis of the single cell transcriptomes revealed that tumor-infiltrating CAR T cells were clearly distinguished from those circulating in blood (Figure 6.B). CAR T cells separated further into CD4^+^ and CD8^+^ populations (Figure 6.C). In blood, cell cycle stage inference from RNA expression profiles revealed a large stationary population and a small proliferating population of CAR T cells (Figure 6.D). Compared to circulating cells, CAR T cells from tumor were generally in a more activated, proliferative state, with a gradient-like separation of predicted cell cycle phases (Figure 6.D).

**Figure 6.**
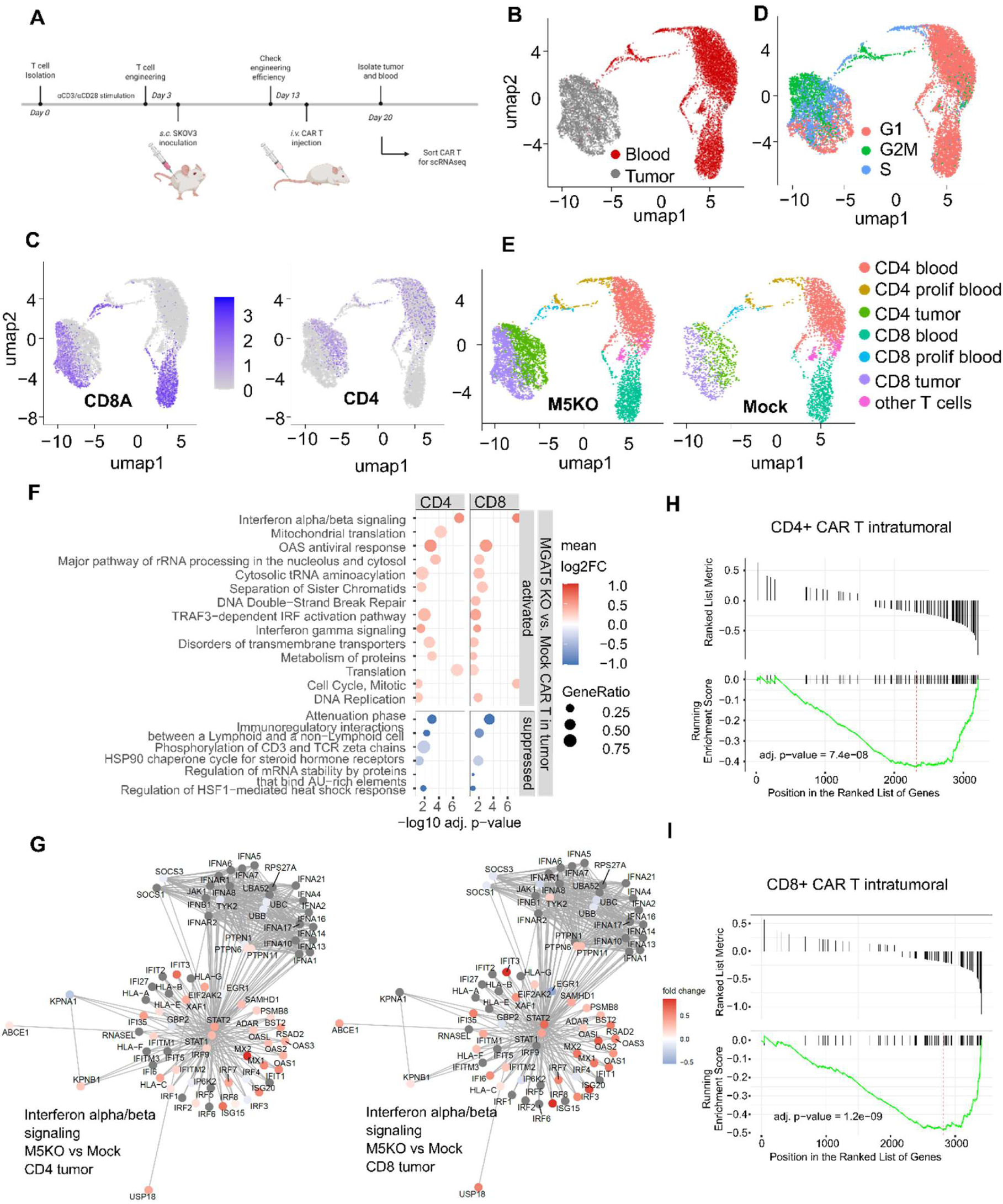
Single cell RNA sequencing of tumor-infiltrating and circulating CAR T cells. A. Experiment outline, performed on donor J. Dimensionality reduction and clustering of cells was driven by key biological differences. B. Sample origin, C. lymphocyte type and D. cell cycle state, leading to the distinction of 7 cell populations as depicted in panel E. F. Gene set enrichment analysis of Reactome pathways and (G) significantly upregulated interferon alpha/beta signaling, compared to control CAR T cells. Genes not included in differential expression analysis (expressed in fewer than 25% of cells or showing log2 fold change < 0.1) were marked in grey H-I. Gene set enrichment analysis showed significant overlap between our results (CD4 tumor, panel H; CD8 tumor, panel I) and previously published genes implicated in CAR T cell exhaustion^42^. The running enrichment score is calculated by traversing our ranked list of genes, increasing a running sum when a gene from the tested set is encountered and decreasing when a gene outside the set is encountered. The enrichment score (dashed red line) is the maximum deviation of this running sum from zero.

Based on CD4/CD8 state, blood/intratumoral location and inferred cell cycle phases, we defined 7 clusters (Figure 6.E) for further exploration of transcriptional differences associated with MGAT5 KO. Subsequently, differential expression (DE) analysis was performed within the same cluster, revealing a total of 193 significant DE genes between MGAT5 KO and non-glycoengineered in at least one of the clusters (Wilcoxon Rank Sum test for genes expressed in at least 25% of cells of one of the compared populations and minimal fold change threshold of 0.1, Bonferroni-corrected p-value of at least 0.05; Supplementary Table 1 We also performed gene set enrichment analysis of Reactome pathways to perform the differential analysis at functional pathway/module level (Supplementary Table 2), visualized according to statistical significance level, fraction of pathway genes differentially regulated and mean log2 fold change of gene expression of pathway genes in Figure 6.F^40^. The Reactome pathways related to active cell cycle progression were more prominently upregulated in MGAT5 KO cells, whereas for the CD4^+^ subfraction, we observed more significant upregulation in MGAT5 KO cells of ribosomal biogenesis and translation, a known hallmark of CD4^+^ cell activation^41^ (Figure 6.F). A somewhat higher proportion of active cell cycling (in G2M and S phase) was observed in both the intratumoral CD4^+^ and CD8^+^ subfraction (Supplementary Figure 7.A). Furthermore, across both CD4^+^ and CD8^+^ intratumoral fractions, strikingly, pathways related to IFN type I signaling were significantly more activated upon MGAT5 KO in both CD4^+^ and CD8^+^ CAR T cells (Figure 6.F and G), a feature of persistently activated T cells which we discuss further below as it is a topic of significant current interest in CAR T cell enhancement-engineering. In blood, transcriptional differences were minor between MGAT5 KO and non-glycoengineered cells, barely reaching statistical significance (Supplementary Figure 7.B). The overall picture that emerges is that of MGAT5 KO CD70 nanoCAR T cells more vigorously mounting a sustained proliferative cytotoxic antitumoral response through both a CD4^+^ and a CD8^+^ T cell contribution, so it was interesting to compare our data with the transcriptional signature previously reported for human CAR T cell exhaustion by prolonged antigen exposure^42^. Remarkably, MGAT5 KO indeed significantly suppressed the gene signature associated with CAR T cell exhaustion by long antigen exposure (Figure 6.H and I).

## Discussion

The engineering of glycosylation in order to enhance biopharmaceutical functionality has met with previous successes. The most notable example in the area of cancer immunotherapy is the discovery that antibody dependent cellular toxicity potency of hIgG monoclonal antibodies could be boosted by engineering of the production cell lines in various ways to reduce or abolish core-α-1,6-fucosylation of the conserved CH2-domain linked N-glycan^43–45^. The strongly enhanced affinity to FcγRIIIa achieved in this way is now at the basis of multiple enhanced anti-tumor antibody therapies^46–49^. The mammalian cell glycocalyx constitutes a complex array of precisely regulated carbohydrate structures on transmembrane and secreted proteins as well as on plasma membrane glycolipids. These glycans are biosynthesized in the cell’s secretory system by an array of glycosyltransferases and glycosidases. Cellular differentiation pathways as well as cellular physiology alterations often result in cell-type and cell-state-specific glycocalyx composition differences. This modulates altered interactions with the cellular environment, resulting in more or less cell motility, adhesion, shaping of growth factor gradients, signaling receptor clustering at the cell surface etc. In principle, this should offer novel biopharmaceutical opportunities for altering complex behavior of therapeutic cells. This was the rationale for us to systematically start exploring the ‘deconstruction’ of the CAR T cell glycocalyx through genome editing of glycosyltransferases that are at key positions in the glycan biosynthetic pathways, and we here report on the first therapeutically relevant results from these ongoing studies.

Poly-N-acetyllactosamine repeat structures occur on both N-and O-glycans of glycoproteins and on certain glycosphingolipids. In human T cells, a large fraction is present on N-glycans, as staining with a lectin for poly-LacNAc was reduced by >50% in our MGAT5 KO cells, which is why we prioritized studying N-glycan poly-LacNAcylation for our studies. As reviewed in the introduction, basic immunology studies over the past few decades, mainly in the mouse, also provided incentive to explore whether MGAT5 inactivation would result in enhanced CAR T cell efficacy. However, promising basic immunology research findings on T cell function in murine model cells do not necessarily translate to technologically useful interventions in CAR T therapy. Hence, to ensure translational relevance of our results, we decided to stick as close as possible to current human CAR T cell manufacturing methods for our studies. We coped with the well-known large variance in quality of CAR T cells produced from different individuals by replicating the laborious tumor control studies in mice with up to 5 human donors per tumor model. We used a highly immunosuppressive human xenograft carcinoma model (SKOV-3) and a very aggressive model of metastatic B cell lymphoma (Raji) to truly put our engineered CAR T cells to challenging tests that reflect the 2^nd^ or 3^rd^ line tumor treatment setting in which CAR T cells are mostly used.

In summary, our data demonstrate that MGAT5 KO has the potential to enhance control of both carcinoma and lymphoma tumors, through sustained elevated tumor cell-killing functionality and proliferation. Especially when CAR T cells of human donors showed poor tumor control, MGAT5 KO boosted this substantially and consistently. This was hallmarked by higher levels of active tumor cell-killing competent CAR T cells in the systemic circulation, persistently constituting effective antitumor ‘immune memory’ that could control secondary tumor challenges. These easy to monitor enhanced blood counts of MGAT5 KO CD70 nanoCAR T cells, especially ∼21 days post CAR T treatment, could constitute a convenient surrogate biomarker for use in translation of our findings to clinical studies. MGAT5 KO does not enhance CAR T cell antigen sensitivity (which may be in contrast to MGAT5 KO effect on TCR-mediated antigen recognition), which we consider as favorable from a safety perspective, as many tumor-associated antigens are also expressed at lower levels on healthy cells^16,20^. Also supporting the safety profile of the MGAT5 KO intervention, the enhanced antigen recognition-triggered proliferation of MGAT5 KO cells is not because of independence from cytokine growth factors, as the cells do not grow at all when these growth factors are withdrawn.

The mild but comprehensive upregulation of type I interferon stimulated genes in MGAT5 KO CD70 nanoCAR T cells infiltrating the tumor suggests enhanced immune surveillance activity, a more inflammatory or cytotoxic phenotype and could potentially be linked to persistent functional activation^50–53^. Comparison of our single cell dataset of intratumoral CAR T cells to published data from a study evaluating the response of T cells to continuous antigen exposure-triggered exhaustion, shows that the gene expression signature linked to exhaustion is indeed suppressed in MGAT5 KO CD70 nanoCAR T cells^42^. Whereas type I interferon signaling is critical in activating and polarizing acute immune responses (most studied in antiviral immunity), strong prolonged signaling dampens T cell activity upon prolonged antigen stimulation such as in chronic viral infections^54,55^. Given this duality, how to intervene optimally in type I interferon signaling to enhance CAR T therapy is not unequivocal in the literature^56–59^. Most studies that completely inactivate type I interferon signaling have observed a level of enhanced CAR T potency, but a very recent preprint publication reveals that strength of the type I interferon signal matters: with low but not high concentrations of interferon-alpha addition during CAR T cell expansion, a more efficacious tumor control was achieved^60^. We propose that our findings on mild type I interferon activation in intratumoral MGAT5 KO CD70 nanoCAR T cells are consistent with this study, usefully achieving this phenotype after the CAR T cells arrive in the tumor, when they are required to persistently activate in order to successfully control the tumor.

Ensuring the technical feasibility of translating our findings to clinical CAR T production, we have implemented MGAT5 KO by adding a CRISPR-Cas9 ribonucleoprotein nucleofection step to the standard workflow that is already used for CAR T manufacturing in the clinic^61–63^, and other preferred technologies could likely be used for this as well^64–67^. It should hence be straightforward to plug in the MGAT5 KO module onto most of the production workflows for almost any CAR T product.

Overcoming the poor proliferation and persistence of CAR T cells during solid tumor therapy, which results in therapeutic failure, is a key and very active focus of research at the present time. Other promising recently reported approaches involve installing autocrine cytokine production in the cells or ablating immune signaling-dampening factors, in particular CD5^68–70^. We propose that CAR T cell surface poly-LacNAc density reduction through inactivation of MGAT5 glycosyltransferase offers a novel highly promising engineering module that targets the glycocalyx of the cells, a very different aspect of T cell biology that remained unexplored as a CAR T engineering target until now.

## Materials and methods

### Ethical approval

Research work with anonymized buffy coats has been approved by the Red Cross Flanders (RKOV_19015, G20240913A). All experiments were approved and performed according to the guidelines of the ethical committee on Medical Ethics of Ghent University, Belgium (EC UZG 2019/1083). We make use of a biobank (Unca-hT cell biobank) which was registered at FAGG (reference number at Bioresource center Ghent is BR-109). The breeding of NSG mice is covered by file E-726 and animal experiments are covered by files EC2020-009, EC2021-063 and EC2022-006.

### Cell lines

We selected three tumor cell lines to study the anti-tumor functionality of the MGAT5 KO CD70 nanoCAR T cells. THP-1 cells are a M4 subtype acute myeloid leukemia (AML) cell line^71^, SKOV-3 cells are a highly immunosuppressive human serous adenocarcinoma cell line^33,34,36,72^ and Raji cells are a human B lymphoblastoid cell line derived from a patient with Burkitt’s lymphoma. We confirmed the cell surface CD70 expression on these cells by flow cytometry (Supplementary Figure 1.A-B). Further, we confirmed absence of CD70 expression on non-transduced (NTC) and CD70 nanoCAR transduced CD3^+^ T cells. THP-1 cells were cultured in RPMI medium (Gibco) supplemented with 10% fetal calf serum (FCS), 0.03% L-Gln, 0.4 mM sodium pyruvate and 50 µM β-mercaptoethanol. SKOV-3 cells expressing luciferase were kindly provided by Dr. Olivier De Wever^73^ (Ghent University, Faculty of Medicine and Health Sciences) and were cultured in DMEM medium (Gibco) supplemented with 10% FCS and 1% penicillin/streptomycin (Pen/Strep). Jurkat cells were obtained from ATCC and were cultured in RPMI medium (Gibco) supplemented with 10% FCS, 2mM L-Gln and 0.4 mM sodium pyruvate. Raji cells expressing luciferase were generated at our labs and were cultured in RPMI medium (Gibco) supplemented with 10% FCS, 2mM L-Gln, 0.4 mM sodium pyruvate, 50 µM β-mercaptoethanol and 1% Pen/Strep. All cell lines were maintained in a 37°C, 5% CO_2_, fully humidified incubator and passaged twice weekly. SKOV-3 cells are detached with trypsin/EDTA and always seeded at 1 x 10^6^ cells in a T175 falcon. Suspension cells were subcultured to 3.10^5^ cells/mL (THP-1/Jurkat) or 5.10^5^ cells/mL (Raji).

### Human CD3^+^ T cell isolation and culturing

Leukocyte-enriched buffy coat samples were obtained from healthy donors attending the Red Cross center after informed consent and ethical committee approval. Peripheral blood lymphocytes were prepared by Ficoll-Paque density centrifugation as described in the instruction manual for Leucosep^TM^ (Greiner bio-one). CD3^+^ T cells were isolated by negative selection with antibodies against CD14, CD15, CD16, CD19, CD36, CD56, CD123 and CD235 (MojoSort^TM^ Human CD3 T cell selection kit, Biolegend) according to the manufacturer’s protocol. Cells were cultured in IMDM + Glutamax medium (Gibco-BRL) supplemented with 10% heat-inactivated FCS and stimulated with Immunocult^TM^ Human CD3/CD28 T cell Activator (Stemcell Technologies) (25 µL/ 10^6^ cells) for 3 days at 37°C in the presence of 10 ng/ mL IL-12 (Biolegend).

Prior to cell seeding, cells were washed twice with PBS before putting them in culture with recombinant human IL-7 at 10ng/mL (Miltenyi) and recombinant human IL-15 at 10 ng/mL (Miltenyi). Cytokines and medium were replaced every 2-3 days. Cell densities were maintained between 1 x 10^6^ and 3 x10^6^ cells/ mL. In experiment with donors A, B, D, E, F, G, H, I, J and K, CD3^+^ isolation was started from freshly prepared PBMC cells while in the experiment with donor C, CD3^+^ T cell isolation was started from a frozen batch of PBMCs (Supplementary Table 3).

### Generating *MGAT5* KO nanoCAR T cells

We first selected a CRISPR-Cas9 guide RNA (gRNA) to delete *MGAT5* in CAR T cells. We designed four gRNAs using the Synthego design tool (https://www.synthego.com/products/bioinformatics/crispr-design-tool). Guides were ordered as chemically modified synthetic sgRNAs (with 2’O-Methyl at three first and three last bases and 3’ phosphorothioate bonds between first three and last two bases) and reconstituted at 100 µM in TE buffer. After initial screening, we selected the most efficient sgRNA (Supplementary Figure 2) on the basis of Sanger sequencing and evaluation of lectin staining as described below. We used this sgRNA to knockout *MGAT5* in primary T cells using an optimized CAR T manufacturing process in which CD3^+^ T cells were stimulated with Immunocult for three days, after which activated T cells were first subjected to Cas9 RNP nucleofection, followed by a retroviral transduction on the same day. Engineering efficiencies were assessed on day 10. The experimental timeline is depicted in Figure 1.B.

Recombinant Cas9-GFP protein was purchased from the VIB protein core (https://vib.be/labs/vib-protein-core). Cas9 RNP was made by incubating Cas9 protein with sgRNA at a molar ratio of 1:2 at 37°C for 15 min immediately prior to electroporation in T cells. Electroporation was performed using the Lonza Amaxa 4D Nucleofector X unit (Program EH-115) and the P3 primary cell kit with the following conditions: 1 x 10^6^ cells/20 µL P3 buffer per cuvette (16-well strips) with 20 µM Cas9-RNP. Following nucleofection, 80 µL pre-warmed medium was added per well and cells were allowed to rest for 30 mins at 37°C, 5% CO_2_, before they were collected in prewarmed medium, centrifuged for 5 min at 300 x g, counted and seeded on retronectin-coated plates for transduction.

0.1 x 10^6^ cells were collected and lysed in QuickExtract^TM^ (Lucigen Epicentre) according to the supplier’s instructions. The target site was amplified by PCR (Supplementary Figure 2) and Sanger Sequenced. Sequencing data was analyzed by variant deconvolution modelling with the ICE tool (Inference of CRISPR Edits, Synthego) to infer the percentage of insertions and deletions (INDEL score) and the percentage of insertions and deletions that are out of frame (KO score)^74^. Cutting and error-prone repair usually results in mixed sequencing bases after the cut. %INDEL: % insertions/deletions, %KO: proportion of indels that indicate a frameshift or are 21+ bp in length (assumes all edits are in a coding region), R2: model fit (how well the proposed indel distribution fits the Sanger sequence data of the edited sample).

### Production of retroviral vectors

In our work, we made use of a nanoCAR targeting CD70, an emerging target across several carcinomas and B cell lymphoma, allowing to use the same nanoCAR design in xenograft models of both very different tumor types. The nanobody used in this nanoCAR construct was identified previously at our center (Dr. Jan Tavernier, VIB-UGent Center for Medical Biotechnology) and was built into standard 2^nd^-generation CAR signaling domain architecture as described previously^26^. The retroviral construct encoding the nanoCAR sequence was previously cloned in the LZRS-IRES-eGFP vector^26^. Viral particles were produced using standard Ca_3_(PO_4_)_2_ transfection of the Phoenix ampho packaging cell line^75^. Retroviral supernatant was collected at day 14 after transfection and puromycin selection and kept at −80°C until use.

### Generation of CD70 nanoCAR Expressing Human T cells

Immunocult-stimulated human CD3^+^ T cells were retrovirally transduced on Retronectin-coated plates (TaKaRa). 500 µL of cells per well at 0.5 x 10^6^ cells/mL were supplemented with 0.5 mL retroviral supernatant and centrifuged for 90 minutes at 890 g at 32°C. Transduced cells were detected by eGFP fluorescence or by staining with an anti-VHH antibody (iFluor555) directed against the nanobody constituting the extracellular domain of the CAR and analyzed by flow cytometry. A minimum of 50 000 events was recorded.

### Lectin-based flow cytometry

For the evaluation of the poly-LacNAc density on the cell surface, we used the lectin from *Datura stramonium* and *Phaseolus Vulgaris* Leucoagglutinin (PHA-L), both at a staining concentration of 10 µg/ mL (Vector laboratories, Biotinylated DSL: B-1185, Biotinylated PHA-L: B-1115). 2 x 10^5^ cells per condition were collected and rinsed three times with PBS. Cells were incubated with fixable viability dye eFl780 (eBioscience, 1000x dilution) and incubated for 30 minutes at 4°C in the dark. After washing the cells with PBS, the cells were fixed with 4% paraformaldehyde (PFA), for 20 minutes at room temperature in the dark. After rinsing with lectin binding buffer (LBB; PBS with 0.1 mM CaCl_2_), cells were stained with biotinylated lectin in LBB for 1 hour at 4°C. After this lectin staining, cells were washed in LBB before adding the PE-coupled neutravidin (Invitrogen, 5 µg/mL) for 30 minutes at 4°C. After washing the cells with PBS, samples were resuspended in LBB and analyzed by flow cytometry. A minimum of 50 000 events was recorded.

### PNGaseF digest and N-glycan analysis using CGE-LIF profiling

In order to prepare cell surface N-glycans for CE-LIF profiling, 1 x 10^6^ cells were collected per condition and washed three times with PBS to reduce the presence of medium-derived glycans. Cell culture medium was collected for N-glycan labeling. PNGaseF digest (0.125 IU/ 1×10^6^ cells, in-house production) was performed in 25 µL final volume in PBS for 2 hours at 37°C. Cells were removed by centrifugation (5 min at 300 x g) and the supernatant was subjected to another centrifugation step (15 min at 15 000 rpm) to remove cell debris. The remaining liquid portion of the sample was stored at −20°C until 8-aminopyrene-1,3,6-trisulfonic acid (APTS) labeling and CE-LIF analysis.

The remaining N-glycan samples were labelled by adding an equal volume (20 µL) of labeling mix consisting of a 1/1 v/v mix of 1M borane-morpholine complex in 20% DMSO, 20% SDS and 4M Urea mixed with 350 mM APTS in 2.4M citric acid and 14% SDS immediately prior to labeling. The labeling reaction was incubated at 70°C for 1 hour and allowed to cool down at 4°C before purification. Size exclusion chromatography (Sephadex G-10 resin with an exclusion limit of 700 Da prepared in a 96-well setup in Multiscreen-Durapore plates) was performed twice to desalt the samples and to remove free unreacted APTS^76^. The labelled glycans were then dried in a speedvac.

Purified labelled and dried N-glycans were resuspended in 10 µL of ultrapure water and analyzed with capillary gel electrophoresis on an eight-capillary DNA sequencer (Applied Biosystems 3500 Genetic analyzer). An internal standard (GlyXera) was added to the samples to be able to align profiles from different samples. Samples were injected on a 50 cm capillary at 15 kV for 10 seconds, using POP7 polymer and 100 mM TAPS, pH 8.0, containing 1 mM EDTA as the running buffer. N-glycan profiles were analyzed through the Genemapper 6 software.

### Flow cytometry analysis

Flow cytometry analysis was performed on 0.2 x 10^6^ cells per sample collected in a 96-well U bottom plate. Cells were rinsed with FACS buffer (PBS containing 0.5% BSA and 2mM EDTA) for 5 min at 300 x g and incubated with Fc Receptor Blocking solution (Human TruStain FcX^TM^, Biolegend) for 10 minutes prior to cell surface staining with fluorescently labelled antibodies in Brilliant Stain buffer (BD Biosciences) for 30 minutes at 4°C.

For human CD3^+^ T cell phenotyping, cells were labelled with fluorescent antibodies against human CD8, CD62L and CD45RA (BD Biosciences) and CD3, CD4, CD25, CD69, and CD279 (PD-1) (Biolegend). A Fixable dye eFluor^TM^ 780 (eBioscience) was used to evaluate live/dead cells.

Flow cytometer calibration was performed using CS&T beads (BD Biosciences). The gating strategy was set based on fluorescence minus one (FMO) controls and retained for all samples. Jurkat, THP-1, SKOV-3 and Raji cell lines and primary human T cells were labelled with fluorescent antibody against human CD70 (PerCP-Cy5.5 labelled) or isotype control (Biolegend) to verify antigen expression as described before^77^. In all analyses, following doublet exclusion, live cells were identified using a fixable viability dye (Molecular Probes, Life Technologies). Data were acquired on a BD Symphony A5 equipped with five lasers (355, 405, 488, 561, 640nm) (BD Biosciences) and analyzed using FlowJo software (Tree Star, Ashland, OR). Ultra-rainbow fluorescent particles (Spherotech, Cat N° URFP-38-5A) are used for alignment in all channels between different experiments. CountBright^TM^ Absolute counting beads (Molecular Probes, Cat N° 11570066) were used to normalize cell counts. An overview of all flow cytometry antibodies used in this study with details on fluorochrome, dilution, company and catalogue number are given in Supplementary Table 4.

### *In vitro* analysis of cytokine production

Glyco-engineered CD70 nanoCAR T cells were stimulated *in vitro* by co-incubation with THP-1 or SKOV-3 tumor cell lines expressing CD70 in a 96-well plate in duplicate. After 1 hour of co-incubation, BD GolgiPlug (Fisher Scientific, 1:250) was added and after an additional 15 hours of stimulation, the cells were harvested, labelled with fluorescent antibodies against CD3, CD4 and CD8, fixed and permeabilized (Fixation/Permeabilization concentrate – diluent, eBioscience, Cat N° 00-513-43 and 00-5223-56) and labelled for intracellular cytokine levels with fluorescent antibodies against TNF-α (BD Biosciences), IFN-γ and IL-2 (Biolegend). Samples were analyzed by flow cytometry as described above.

### *In vitro* analysis of tumor cell killing

Glyco-engineered CD70 nanoCAR T cells were incubated with 2 x 10^4^ THP-1 cells at different effector/target ratios (0; 0.0015; 0.015 and 0.15) in IMDM medium with Glutamax (Gibco) containing 10% FCS and 1% Pen/Strep. Multiple runs of the same conditions were setup, allowing for flow cytometrical analysis at days 0, 3, 7, 10 and 14. In the wells remaining for analysis at day 10 and 14, 2 x 10^4^ THP-1 cells were added at day 7. Cells were labelled with fluorescent antibodies against CD3, CD4 and CD8 for the analysis of T cells and CD33 for the detection of THP-1 cells.

### Statistical analysis of *in vitro* experiments

To analyze the data of the fractions of cytokine-producing cells upon *MGAT5* glycoengineering (Figure 1.I-J), we considered the control setup in which cells were not stimulated to represent a background signal. We subtracted the mean percentage cytokine positive cells measured in the two technical repeats of non-stimulated cells from the corresponding percentage of cytokine positive cells in the stimulated setups (corresponding means of the setup with the same CAR engineering and donor). For each cytokine, we built an ANOVA model in R^78^, with Stimulation, Donor and Group as main effects (all coded as categorical) and allowing for all two-way interactions. Model terms were tested via F tests. The effect of *MGAT5* engineering, averaged over donors and stimulation strategies, was obtained from each final model using the multcomp package^79^ with sandwich estimators^80,81^.

To analyze the number of mock Cas9 and MGAT5 KO CD70 nanoCAR cells in the THP-1 co-culture experiments (Figure 1.L, M, N, O), we started from flow cytometry-based count data. Since the counts had been normalized using counting beads, they were not necessarily integers so we rounded all to the closest integer. We considered each setup with the same donor, E/T Ratio and type of CAR T cells (i.e. mock Cas9 CD70 nanoCAR or CD70 nanoCAR *MGAT5* KO) as a cluster. Since we had two measurements (technical repeats) at each day and the measurements were performed at day 0, 4, 7, 11 and 14, this means we had 10 measurements in each cluster. Furthermore, we observed a slight rise in the counts of the NTC cells over time in the control setups. We corrected the CAR T cell counts for this background (per cluster and at each timepoint) by subtracting the mean background count from the measurements.

We analyzed the background-corrected counts with a generalized linear mixed model (GLMM) to allow for modeling of the within-cluster correlation over time. Since the data showed considerable overdispersion, the most appropriate choice was to use a negative binomial model (with a log link function). The GLMMadaptive package^82^ allows to fit such a model in R^78^ using adaptive Gaussian quadrature (AGQ). We did not have enough data to fit a random slope model, so we settled for a random intercept model of the following model formula:

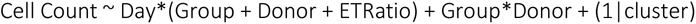

In this model, all predictors (Day, Group, Donor ETRatio) are coded as a categorical variable. We chose to also model the time variable as categorical since the log-transformed counts do not evolve linearly with time. Contrasts were estimated using the multcomp package^79^.

### *In vivo* analysis of glyco-engineered CD70 nanoCAR T cell tumor control efficacy in the SKOV-3 model

NSG mice (breeder pairs obtained from The Jackson Laboratory, breeding in house) between 8-12 weeks of age were subcutaneously (in the right flank) injected with 2 x 10^6^ luciferase-expressing SKOV-3 cells in 50 µL PBS. Ten days after tumor cell inoculation, after establishment of a solid, palpable tumor, mice were treated with 3 x10^6^ of either mock Cas9-engineered or MGAT5 KO CD70 nanoCAR (in 200 µL total volume in PBS). As control groups, mice were treated with PBS to evaluate tumor development, or with non-transduced T cells (NTC) to evaluate graft-versus-host disease (GvHD) and non-specific anti-tumor effects. Body weight and tumor progression was followed up by caliper and BLI. A dose of 150 mg/kg D-luciferin potassium salt (Perkin Elmer) was injected intraperitoneally 10 minutes before BLI. Imaging data were analyzed using Living Image Software and reported as photons/second. Surviving mice were followed up in time and challenged between day 87 and day 90 with a second tumor injection in the left flank (opposite flank of first tumor injection) to mimic a tumor relapse, thus evaluating long-term anti-tumor control efficacy. Again, tumor burden was evaluated over time and the mice were sacrificed between day 118 and day 123 for endpoint analyses (see experimental timeline for details). For statistical analysis, the ‘Untreated’ group contains the data from all the mice that did not receive any CD70 nanoCAR T cells, and thus includes untreated mice and mice treated with PBS or NTC (which all behaved similar with respect to tumor control).

In the experiment with donor A, we started with 10 mice in the PBS and 6 mice in the NTC group and 12 mice in the CD70 nanoCAR mock Cas9/*MGAT5* KO groups. At day 34, 6 mice from the PBS group, 5 mice from the NTC group, 5 mice from the CD70 nanoCAR mock Cas9 and 6 mice of the CD70 nanoCAR *MGAT5* KO group were euthanized and used for analysis of CAR T cell numbers in the spleen and blood. The other 5 mice from the CD70 nanoCAR mock Cas9/*MGAT5* KO groups were kept for tumor rechallenge. In the experiment with donor B, we started with 6 mice in the PBS and NTC group and 9 mice in the CD70 nanoCAR mock Cas9/*MGAT5* KO groups. We also included 9 mice in a CD70 nanoCAR group (i.e. with cells that did not receive mock nucleofection). All mice were kept for rechallenge, unless the pre-defined humane endpoint was reached (which only occurred in control mice that did not receive CAR T treatment). Humane endpoint is reached when mice lost > 20% of their body weight. In the experiment with donor C, we started with 7 mice in the NTC group, 13 mice in the CD70 nanoCAR mock Cas9 group and 14 mice in the CD70 nanoCAR *MGAT5* KO group. All mice were kept for rechallenge, unless humane endpoint was reached (occurring in non-CAR T treated control mice). At the start of each experiment, we also kept a group of 8 (donor A) or 10 (donor B/C) mice to be used as age-matched control (=PBS) group in the tumor rechallenge phase of the experiment.

### *In vivo* analysis of glyco-engineered nanoCAR T cell tumor control efficacies in the Raji model

Female NSG mice (breeder pairs obtained from The Jackson Laboratory, breeding in house) between 8-15 weeks of age were intravenously injected with 0.5 x 10^6^ Raji cells in 50 µL of PBS. 4 x 10^6^ CD70 nanoCAR T cells were intravenously injected 7 days post-Raji injection (in 200 µL total volume in PBS). In preliminary titration experiments using non-engineered CAR T cells from a different donor, we had determined that a dose of 4 x 10^6^ CD70 nanoCAR T cells per mouse yielded a delayed tumor growth in the intravenous Raji model, while not being sufficient for survival, such that improvements in efficacy could be detectable by increased survival rates. Body weight was followed up as was tumor progression by BLI. A dose of 150 mg/kg D-luciferin potassium salt (Perkin Elmer) was injected intraperitoneally 10 minutes before BLI. Imaging data were analyzed using Living Image Software and reported as photons/second. In contrast to the subcutaneous localized SKOV-3 carcinoma model, this Raji lymphoma model is one of hematogenous aggressive metastasis, with tumors rapidly establishing in multiple organs, including the brain.

In the experiment with donor D, we started with 5 mice in all groups. In the experiment with donor E, we started with 4 mice in all groups. In the experiment with donor F, we started with 5 mice in the NTC group, 5 mice in the mock Cas9 nanoCAR group and 4 mice in the MGAT5 KO CD70 nanoCAR groups. In the experiment with donor G, we started with 7 mice in the NTC group, 8 mice in the mock Cas9 CD70 nanoCAR group and 6 mice in the MGAT5 KO CD70 nanoCAR group.

### Endpoint and intermediate analysis on spleen and blood

The presence and phenotype of CAR T cells was evaluated in the blood and spleen. Peripheral blood was collected via the tail vein and transferred to EDTA coated Microvettes (Sarstedt) for intermediate analysis at day 34 (donors A,B,C) and at days 80 (donor A), day 88 (donor B) and day 85 (donor C) in the SKOV-3 model and at day 27 (donor D, E and G), day 31 (donor F), day 95 (donor E), day 94 (donor F) and day 90 (donor G) in the Raji model. At the end of the experiments (see time lines for exact days on Figure 2 and Figure 3), blood was also collected after euthanasia and puncture of the right atrium of the heart. The volume of blood was determined and red blood cells were removed using ammonium-chloride-potassium (ACK) lysis buffer (added in a 1:20 blood:ACK volume ratio; Lonza) followed by neutralization in FACS buffer and centrifugation at 300 x g for 5 min. Cells were counted prior to antibody staining for flow cytometry analysis.

In the SKOV-3 model, spleens were dissected from all of the surviving mice at the end of the experiments, (see time lines for exact days on Figure 2). For donor A, where all of the CAR T treated mice in both non-engineered and MGAT5 KO groups had already cleared the tumor at day 34, half of the mice in the experiment were euthanized at that timepoint for spleen investigation. Spleens were collected and processed to a cell suspension through a 70 µM cell strainer. Cells were counted prior to antibody staining for flow cytometry analysis.

### Statistical analyses *in vivo* experiments

#### FACS-based cell count data from blood and spleen (SKOV-3 model) with donors A, B and C and from blood (Raji model) with donors E, F, G and H

To compare the FACS-based cell-count data between the *MGAT5* KO and WT CAR T groups, we ran a separate statistical model for each setting. For the SKOV-3 model, the settings are comprised of all unique combinations of donor (A, B or C), timepoint (Early, Mid, Late/Endpoint), and tissue source (spleen). Excluding settings for which data was lacking, this resulted in 13 statistical tests (see below) corresponding to the panels in Figure 2.E and F. For the Raji model, the settings are comprised of all unique combinations of donor (E, F, G, H), timepoint (timepoint 1, timepoint 2) and target (CD70). Excluding settings for which data was lacking, this resulted in 13 statistical models (see below), corresponding to the panels in Figure 3.D. The lack of data for some settings of the blood count analysis should be read in the context of survival curves presented in the same figure; in these groups, mice had already been euthanized or died at the time of blood cell count analysis.

We ran quasi Poisson models in R^78^, using the glm function with Cell Counts per mL of blood or number of cells per spleen as the outcome variable and Group (*MGAT5* KO vs WT) as the predictor variable. The presented p-values were then either directly extracted from the glm (Raji model), or calculated from the glm using the multcomp package^79^ (SKOV-3 model).

#### Treatment of tumor-bearing mice with MGAT5 KO CD70 nanoCAR T cells leads to a better control of tumor growth rate

To evaluate differences in tumor growth or resolution between the treated mice, we fitted a generalized linear mixed models (GLMMs). A separate model was fitted for data from each donor and the primary and secondary tumor (Supplementary Figure 4, Supplementary Figure 5, Supplementary Figure 6). The main reason for this is that the model would become unnecessarily complex because the timescales (design) of both experiments differ slightly as do the times at which the mice start to respond to the CAR T cell therapy. The latter is possibly due to inherent differences between the CAR T cell batches (i.e. a donor effect).

#### Longitudinal analyses

Tumor volumes were measured by measuring the length and width of a tumor and using the length* width*width/2 (this is a half cube or cuboid) approximation of the volume of a sphere. We cross-checked with BLI data for the small tumors, since this gives a better indication on whether there actually is still a tumor present or whether the lesion is remnant scar tissue. Whenever a small tumor was measured or a “zero volume” was registered (i.e. no morphological lesion remaining), BLI was used to verify whether tumor cells were actually present or not and the caliper measurements were adapted accordingly: when no tumor was found on BLI, we left the caliper measurements to zero, and when a tumor was found on BLI but not measurable by caliper, we set the tumor volume equivalent to 0.0625 mm^3^ (corresponding to 0.5 mm width/length).

We then analyzed the tumor volume data (in mm^3^) of each experiment (donor A, B and C) and each phase (primary tumor before and after rechallenge and secondary tumor) separately by fitting a GLMM to each using the glmmTMB package^83^ in R^78^. The models used a log link and a Tweedie error structure to accommodate exact zeros and the mean-variance relationship in the data. Further, we used natural splines (with df = 3 to 5) to model the (continuous) time variable in an interaction with the (categorical) group variable. This allows for flexible modeling of the tumor volume in function of time in each treatment group. Random effects included a per-mouse random intercept and, if necessary, a random slope for the time variable to model within-mouse correlation over time. The models were fitted using REML. Contrasts were calculated using the multcomp package^84^ to obtain multiple-testing adjusted p-values and 95% confidence intervals.

#### Survival analysis (time to relapse)

To analyze the time to relapse, we first defined the start of follow up as the moment the primary tumor was controlled or partially controlled. We define control as the first day the tumor became completely undetectable on BLI and by caliper measurement. We define partial control as the first day a tumor (that never fully disappears) stopped increasing in size according to caliper measurements and no increase in tumor volume was seen in at least 4 consecutive measurement days. Next, we define a relapse event as the moment a tumor starts growing again. As the onset of relapse, we take the first day that the tumor volume reaches again > 4 mm^3^ and only when tumor growth is also seen in 4 consecutive measurement days following that day. The time to event is then the time between start of follow up and a relapse event and the follow up time is the time between start of follow up and either an event or the end of follow up in case of no relapse. We used R^78^ with the survival^85,86^ and survminer^87^ packages to generate Kaplan-Meier plots with estimates of the median survival times and a corresponding risk and events table. Since relapses were only observed in experiment with donor B, we ran a straightforward analysis with group as the only predictor (groups: CD70 nanoCAR or CD70 nanoCAR -MGAT5 KO). We tested for the difference in survival probability in these groups with a logrank test as implemented in the survival package.

#### *Ex vivo* analysis of tumor cell killing

SKOV-3 tumors in NSG mice were established, treated with CAR T cells and monitored as described above (in this experiment with donor H and I). The number of mice treated were: non-transduced control (NTC) (n=13), CD70 nanoCAR T (n=13), *MGAT5* KO CD70 nanoCAR T (n=9). Peripheral blood was collected via the tail vein and transferred to EDTA coated Microvettes (Sarstedt). The volume of blood was determined and red blood cells were removed using ACK lysis buffer (Lonza) prior to antibody staining for flow cytometry analysis (determination of # circulating CAR T cells in the blood). Each sample was measured separately to determine the amount of circulating CAR T cells. Afterwards, cells were pooled per condition. These blood cells, containing differentially engineered CD70 nanoCAR T cells, were incubated with 2 x 10^4^ THP-1 cells at different effector/target ratios (i.e. cells isolated from 50µL of blood and from 40µL of blood) in IMDM medium with Glutamax (Gibco) containing 10% FCS and 1% Pen/Strep. Cells were labelled with fluorescent antibodies against CD3, CD4 and CD8 for the analysis of T cells and CD33 for the detection of THP-1 cells, at the start of the co-culture (day 0) and at day 4, 8, 11 and 14. At day 8 of co-culture, 2 x 10^4^ THP-1 cells were added to the remaining wells for analysis at day 11 and 14. Cell numbers were determined by flow cytometry.

### CD107a degranulation assay

Membrane expression of CD107a represents a surrogate marker of cytotoxicity of activated and degranulating immune cells. Recombinant Human CD27 Ligand (R&D systems) was coated overnight at different concentrations (0.01 – 10 µg/mL) on the surface of a 96-well plate (Falcon^TM^ clear flat bottom TC, Corning, France) at 4°C. Before the addition of the cells/anti-CD107a antibody, the recombinant human CD27 ligand was removed from the plates and the wells were washed 1x with PBS. CAR T cells (from donor K) were put at 1 x 10^6^ cells/mL (corrected based on %GFP-positivity) and mixed with anti-CD107a-BUV395 antibody at a 10:1 ratio. 50µL of this mix was added per well of the coated 96-well plate. In the control condition, 1,25µL of Immunocult^TM^ Human CD3/CD28 T cell Activator was added (25 µL/ 10^6^ cells). Plates were incubated for 1 hour at 37°C C and 5% CO_2_, and for an additional 4 h in the presence of the secretion inhibitor Brefeldin A (Cayman Chemical).

We analyzed the results for CD107a staining on CAR T cells with anti-CD107a-BUV395 (BD Biosciences) and anti-CD3-BV510 (Miltenyi Biotec) antibodies, in combination with eFluor780 viability dye (Ebioscience) on GFP-positive cells using flow cytometry (BD Symphony A5).

### Cytokine Independent growth assay

Human CD3^+^ T cells were isolated and engineered as described (from donor G). Ten days after engineering and cultivation with IL-7/IL-15, engineered CAR T cells were cultured in IMDM + Glutamax medium (Gibco-BRL) supplemented with 10% heat-inactivated FCS with or without supplemental cytokines. All cells were maintained at a cell density of <1.0 x 10^6^ cells/mL over the course of the experiment. Total cell counts of cells cultured in IMDM with or without supplemental cytokines was monitored for 14 days. Cells cultured in IMDM with supplemental cytokines were given 10 ng/mL of IL-7 and IL-15 on day 0 and subsequently maintained with IMDM containing IL-7 and IL-15 until day 14.

Single cell RNA sequencing (scRNAseq)

### Preparation cells from tumor

SKOV-3 tumor inoculation and CD70 nanoCAR T treatment was performed as described above. Cells from donor J were used. Tumors were collected in PBS and transferred into a gentleMACS C tube (Miltenyi) containing digestion buffer (MEMα (Capricorn), 1% Pen/strep (Sigma), 50 µM β-mercaptoethanol (Gibco), 0,1 mg/mL DNAse I (Roche), 0.125 mg/mL Collagenase D (Roche), 0,1 mg/mL dispase II (Sigma)), GentleMACS C tubes are loaded in the MACS dissociator without heaters and progam h_cord1 was run. The tubes were subsequently incubated at 37°C for 40 min, and vortexed every 10 minutes. After that, the tubes were put on ice and 10mL of ice cold FACS buffer is added to the tube, before passing the suspension through a 70µm cell strainer. The strainer is washed with 10mL FACS buffer and cells are pelleted at 300 x g, 5 min at 4°C. Cells are resuspended in FACS buffer before proceeding with staining.

### Single cell library preparation and sequencing

Mice were sacrificed seven days post nanoCAR T cell injection (from donor J). Peripheral blood was taken following severing of the right atrium of the heart and transferred to EDTA coated Microvettes (Sarstedt). The volume of blood was determined and red blood cells were removed using ACK lysis buffer (Lonza) prior to antibody staining. Tumors were isolated and processed to obtain a single cell suspension as described above. Single cell suspensions from 6 mice/condition were pooled and labelled with anti-human CD45-PerCP-Cy5.5 antibody (BD Biosciences) and a unique human TotalSeq-C hashing antibody (Biolegend) diluted 1:250. To enable multiplexing, we applied a cellular hashing strategy using TotalSeq-C anti-human hashtag antibodies specific against human CD298 and B2M (BioLegend). Each sample received a distinct TotalSeq-C barcode, allowing downstream demultiplexing and assignment of single-cell transcriptomic data to individual samples/conditions. FACS-sorted CD45^+^ single-cell suspensions were resuspended at an estimated final concentration of 1000 cells/μL, and loaded on a Chromium GemCode Single Cell Instrument (10× Genomics) to generate single-cell Gel beads-in-EMulsion (NextGEM). The cDNA libraries were prepared using the GemCode Single Cell 5′ v3 Gel Bead and Library kit, version 5.2 according to the manufacturer’s instructions (10x Genomics). Size selection with SPRIselect Reagent Kit (Beckman Coulter, B23318) was used to separate amplified cDNA molecules for 5′ gene expression and cell surface protein construction libraries. The cDNA content of pre-fragmentation and post-sample index PCR samples was analyzed using the Fragment Analyzer (Agilent). Sequencing libraries were loaded on an Illumina NovaSeq flow cell at VIB Nucleomics core on the Illumina NovaSeq 6000 platform with a read configuration of 28 cycles for Read 1 (which reads the 16bp 10x GEM-cell barcode followed by the 10bp UMI), 10 cycles for i7 index (which reads one of the sample barcodes), 10 cycles for i5 index (which is used for dual indexing), and 90 cycles for Read 2 (which reads the cDNA and de feature barcode (TotalSeq/antibody tags). 1% PhiX DNA was incorporated as a sequencing control, pooled in a 90:10 ratio for the combined 5′ gene expression and cell surface protein samples, respectively.

### Processing of scRNA-seq and hashtag data

The Cell Ranger pipeline (10× Genomics, version 7.2.0) was used to perform sample demultiplexing and to generate FASTQ files for read 1, read 2, and the i5, i7 sample index for the gene expression protein libraries. Read 2 of the gene expression libraries was mapped to the reference genome GRCh38. Subsequent barcode processing, unique molecular identifiers, filtering, and gene counting was performed using the Cell Ranger suite (10× Genomics). Downstream analyses were performed with Seurat^88^ (version 5.0.0). In order to maintain explicit control over all gene and cell quality control filters, we used the raw feature-barcode matrix generated by CellRanger. Raw counts of the samples were normalized using global-scaling normalization and log-transform method using the Seurat NormalizeData function with default parameters. Feature counts for each cell are divided by the total counts for that cell and multiplied by the scale.factor = 10,000 followed by natural-log transformation using log1p. Initial dimensionality reduction and clustering was performed using the Seurat functions FindVariableFeatures, RunPCA, FindNeighbors and FindClusters, with default parameters. Hashtag demultiplexing was performed using Seurat’s HTODemux and MultiSeq and we required agreement of both algorithms for the assignment of cells to their respective biological samples. Cells with inconsistent demultiplexing calls, as well as cells identified as negative and double were removed. Next, we excluded all cells with detection of transcripts of less than 200 genes, as well as genes expressed in less than 3 cells. Subsequently, clusters of mouse cells were identified and removed based on overrepresentation of reads mapping to the mouse genome. After excluding incorrectly demultiplexed cells and mouse cells, we obtained data on 12039 cells across four samples under analysis (mock engineered versus MGAT5 KO CD70 nanoCAR T cells, circulating in blood vs tumor-infiltrating). We repeated dimensionality reduction and clustering on the Seurat functions FindVariableFeatures, RunPCA, FindNeighbors and FindClusters, using the 2000 most variable genes, 12 first principal components and cluster resolution of 0.5. Unsupervised Uniform Manifold Approximation and Projection (UMAP) analyses implemented in the Seurat pipeline were further used to identify the major cell populations. Tumor-infiltrating CAR T cells clustered separately from circulating CAR T cells. For improved discrimination of CD4 and CD8 cells, clustering was further refined using the FindSubCluster function using resolution up to 0.2 and manual curation. CellCycleScoring function was used to distinguish cells in different cell cycle phases. Finally, cell clusters were labelled based on their association with biological samples, CD4 and CD8 expression and proliferation state. Differential gene expression analysis between conditions within the same cell cluster was performed using FindMarkers function with Wilcoxon Rank Sum test for genes expressed in at least 25% cells of one of the compared populations and minimal fold change threshold of 0.1, using the Bonferroni-corrected p-value of at least 0.05. Gene identifier conversions were performed using biomaRt version 2.58.2 and human Ensembl database version 113 (Oct 2024). Comparison to custom gene sets of differentially expressed proteins from other studies was performed using the gene set enrichment function of the clusterProfiler package^89^ (version 4.10.1). Enrichment analyses of pathways from KEGG, WikiPathways and Reactome databases were performed using packages clusterProfiler (version 4.10.1), ReactomePA^90^ (version 1.46.0) and enrichR^91^ (version 3.2) with Benjamini-Hochberg adjusted p-value cutoff of 0.1. Reactome pathways were further narrowed down based on the presence of significant DE genes and average gene log2FC. Shared genes were additionally used to remove redundancy between pathways. Full results are available is Supplementary Table 2

### Amnis ImagestreamX Mark II data acquisition

Samples were analyzed on a Cytek® Amnis® ImagestreamX MkII imaging flow cytometer using Cytek Amnis INSPIRE software version 6.2 (Merck Millipore, Nottingham UK). Prior to experimental analysis, following the manufacturer’s instructions for appropriate ImageStream X Mark II® set up and quality control procedures, system calibrations were performed using ImageStream X Mark II® SpeedBead calibration reagents. 100 µL aliquots of cell suspensions (SKOV-3, THP-1, Raji) at a concentration of∼ 2 × 10^7^ cells/mL in PBS were prepared in 1.5 mL Eppendorfs. Prior to data collection, laser intensities were balanced to limit saturation events and fluorescence channel overspill. For data acquisition an INSPIRE® template was set up for in-focus and single cellular event gating, using Aspect Ratio and Root Mean Square (RMS) features, ensuring collected cells were sufficiently circular and in focus. Data were acquired at a low velocity of 66.0 mm/s, with single cell events being collected at a magnification of× 40. Acquisition of the images for CD70-PerCP-Cy5-5 and Live/Dead eFl780 fluorescence assessment occurred in Channel 5 and 12; bright field: channel 1 and 9; Side scatter: Channel 6. Data analysis was performed using IDEAS® software version 6.0. For SKOV-3 cells 6519 single cells were analyzed, for THP-1 cells: 6208 single cells; for Raji cells: 4830 single cells. Area (µm^2^) and antigen staining intensity for each single event was determined and averages per cell were calculated. Intensity/µm^2^ was plotted.

## Data availability

The single cell RNA sequencing data that support the findings of this study are openly available in the Gene Expression Omnibus (GEO) (GSE301470). Code for the statistical analysis and visualization is made available at GitHub (https://github.com/CallewaertLab/MGAT5KO_CART_manuscript).

## Acknowledgements

Research of EDB was funded through a predoctoral fellowship at FWO, a doctor-assistant mandate at UGent and an IOF Advanced project – F2024/IOF – Advanced/031. MG and LDP were funded through a predoctoral fellowship of FWO. SDM is a junior post-doctoral fellow at FWO supported by the personal grant 12AP724N. This work was further supported by grants G050420N and G028220N of FWO Vlaanderen, a Young Investigator Proof of Concept (YIPOC) grant of the Cancer Research Institute Ghent (CRIG), a Stichting tegen Kanker grant (F/2022/2010) and by the European Union (ERC Advanced Grant, GlycoCAR, Horizon 101098331) as well as by institutional core funding of VIB. We thank Prof. Dr. Y. Chen (Parker Institute for Cancer Immunotherapy Center at UCLA, Los Angeles, CA, USA) for intensive experimental training in the CAR T field and Prof. J. Tavernier for providing the anti-CD70 nanobody used in our CAR designs. We thank the VIB Bioimaging core Ghent (https://vib.be/labs/vib-bioimaging-core-ghent), the VIB Flow Core Ghent (https://vib.be/labs/vib-flow-core-ghent), the VIB Nucleomics Core (https://nucleomicscore.sites.vib.be/en) and the VIB Single Cell core (https://singlecellcore.sites.vib.be/en) for technical support and infrastructure access. We thank Prof. O. De Wever (Ghent University, Faculty of Medicine and Health Sciences) for providing the SKOV-3 cells expressing luciferase. N.C. dedicates this study to the memory of his father Eric Callewaert, who succumbed in the course of our work to one of the tumor types studied here.

## Author contributions

Conceptualization: EDB, NF, NC Methodology: EDB, NF, SDM, NC Software: LM, DF

Validation: EDB, NF, LM, DF, NC Formal analysis: EDB, NF, LM, DF, NC

Investigation: EDB, NF, EPL, LV, AVH, EW, MG, LDP, SVdB Resources: EDB, NF, LM, DF, SDM, BV, NC

Data curation: EDB, NF, LM, DF, NC

Writing – original draft: EDB, NF, LM, DF, NC Writing – review & editing: EDB, NF, NC Visualization: EDB, NF

Supervision: EDB, NF, NC Project administration: EDB, NF

Funding acquisition: EDB, NF, NC

## Supplementary information

**Supplementary Figure 1.**
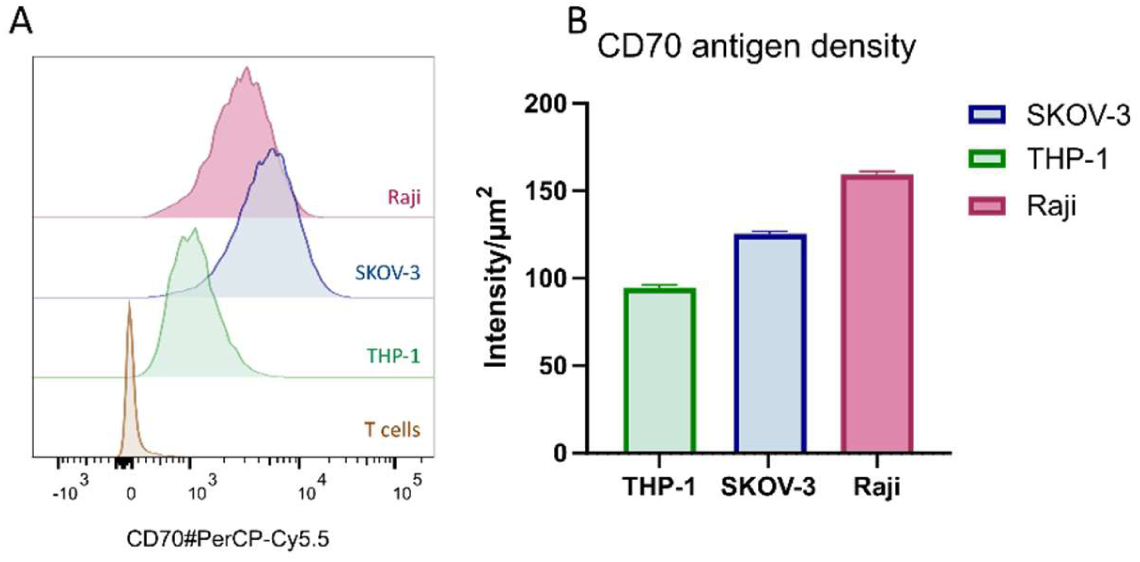
Characterization of the SKOV-3, THP-1 and Raji cell lines used as target cells. A. CD70 antigen expression on the THP-1 and SKOV-3 target cell lines was evaluated by flow cytometry. Jurkat cells were included as negative control. Non-transduced control (NTC) and CD70 nanoCAR expressing human T cells were included to check for auto-antigen expression. B. CD70 antigen density on SKOV-3, THP-1 and Raji target cell lines was evaluated by multispectral imaging flow cytometry. Antigen density is represented as mean fluorescence intensity/ µm^2^. Error bars represent the 95% confidence interval.

**Supplementary Figure 2.**
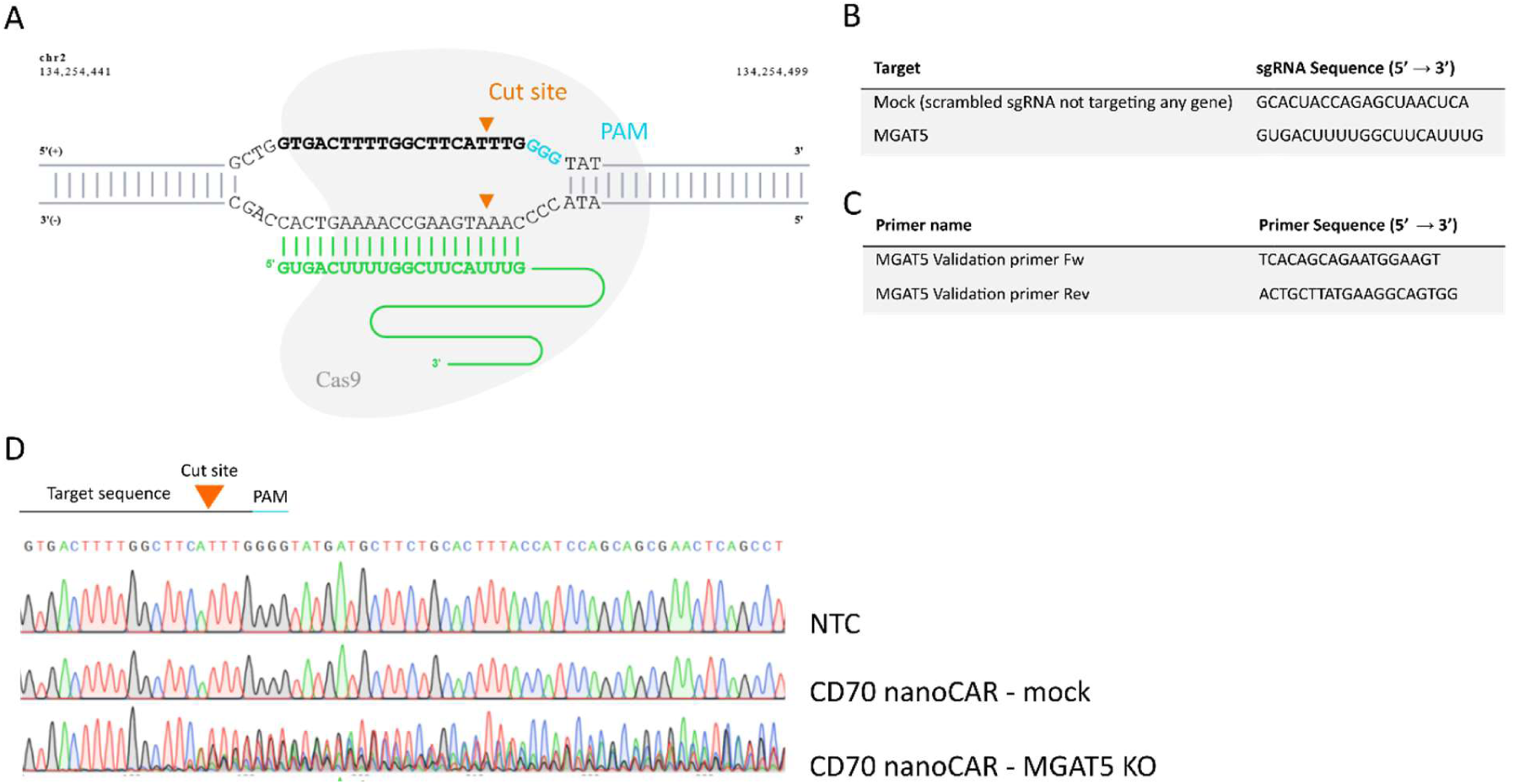
A. Genome localization of the gRNA target site (green) with indication of the PAM (blue) and the predicted Cas9 cut site (orange) B. Overview of the gRNA used in this study to generate a knockout in *MGAT5* C. PCR primers to amplify the CRISPR target site. The forward primers were used for Sanger sequencing. D. Sequencing data of NTC, Mock Cas9 and MGAT5 KO CD70 nanoCAR T cells. *MGAT5*-targeting sgRNA sequence with its PAM and Cut site are indicated in the sequence.

**Supplementary Figure 3.**
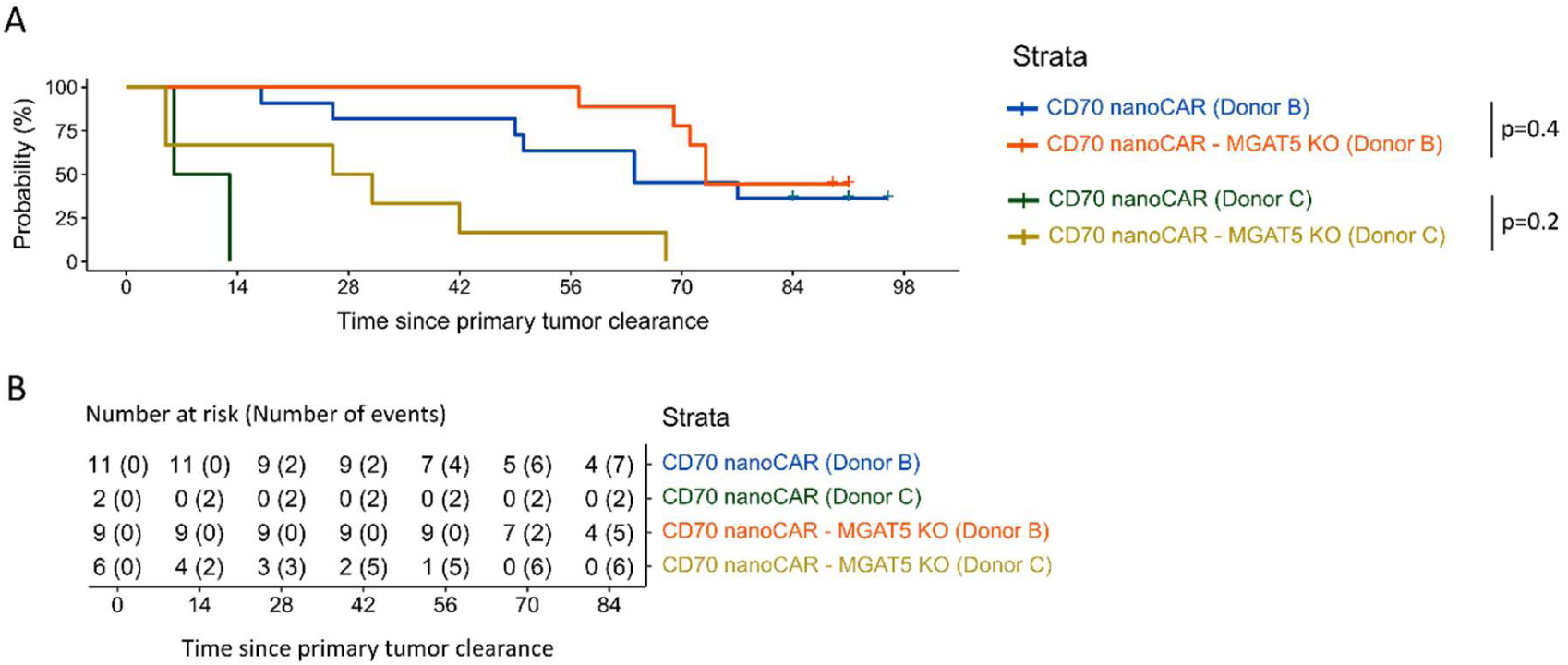
Time (in days) to relapse of the primary tumor (Experiment with donor B/C, not for donor A, as no primary tumor relapse occurred in CAR T treated mice, see Figure 2.B). Time zero was set as the time that the primary tumor was undetectable or partially controlled. Relapse event was defined at the time at which the tumor starts growing again. As the onset of relapse, we take the first day that the tumor volume reaches again > 4 mm^3^ and only when tumor growth is also seen in 4 consecutive measurement days following that day. A. Kaplan-Meier curve. This plot shows the probability of relapse-free survival in the two groups. B. Risk and event table corresponding to the Kaplan-Meier plot. The table shows the number of mice at risk and, between brackets, the cumulative number of relapses in each group and at each time.

**Supplementary Figure 4.**
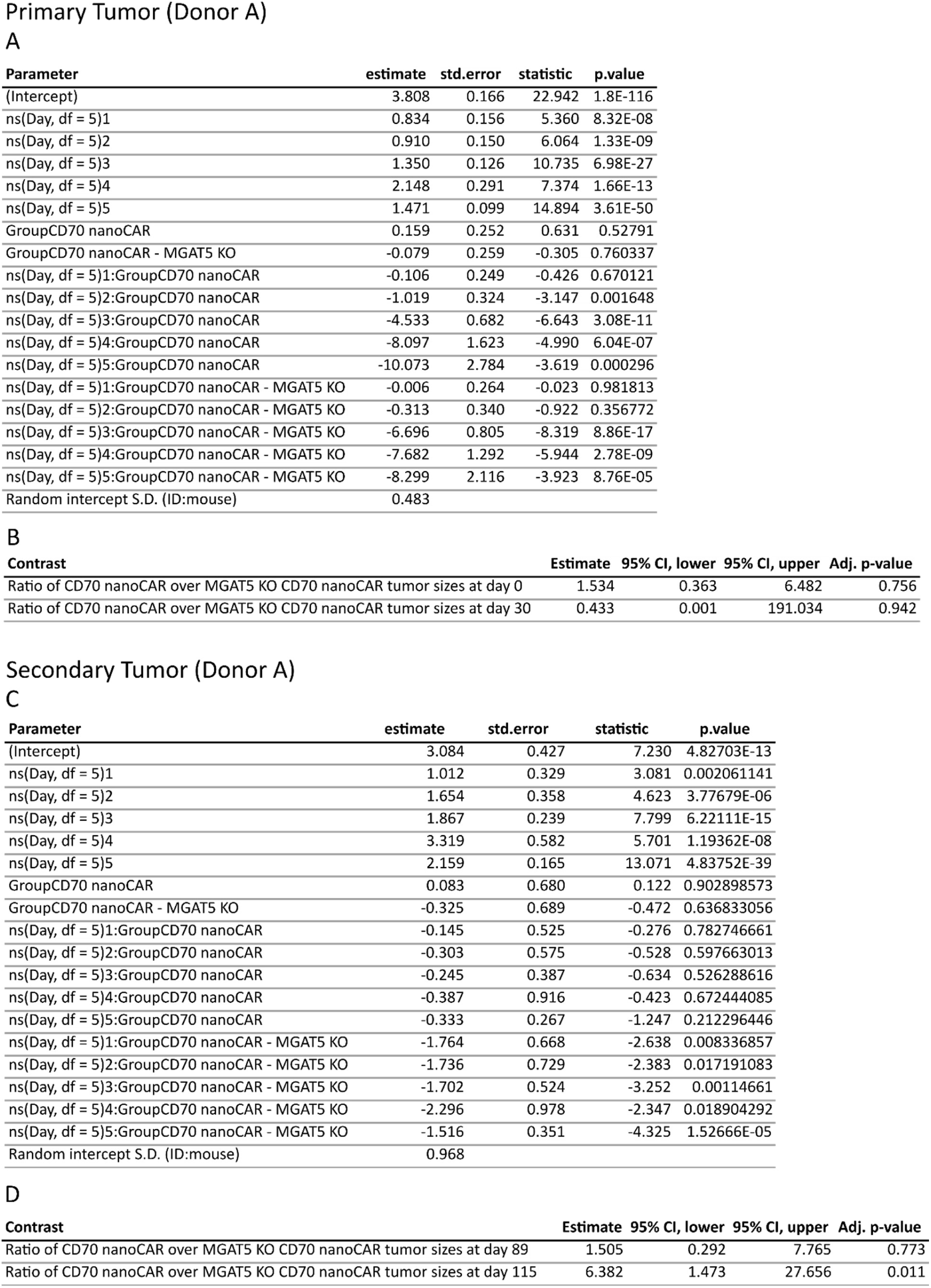
Longitudinal analysis of donor A. Using the longitudinal data of the primary and secondary tumors in experiment with donor A, a Tweedie GLMM (generalized linear mixed model) with log link was used to model the tumor volume of the primary tumor (A) and secondary tumor (C). Summary of the model output, listing all parameter estimates for the model TumorVol ∼ ns(Day, df = 5)*Group + (1 | ID), ns = natural spline. The table gives parameters and standard errors on the natural log scale together with test statistics and multiple testing adjusted p-values (null hypothesis: parameter equal to zero). B and D. Inference for different research questions. In this table, the estimates and confidence intervals are transformed back to the original scale so we can interpret them as the ratio of the mean tumor volume in the WT over the MGAT5 KO group. In this context, the adjusted p-values also relate to a transformed null hypothesis (i.e. estimate equals one). S.E.: Standard Error. CI: Confidence Interval.

**Supplementary Figure 5.**
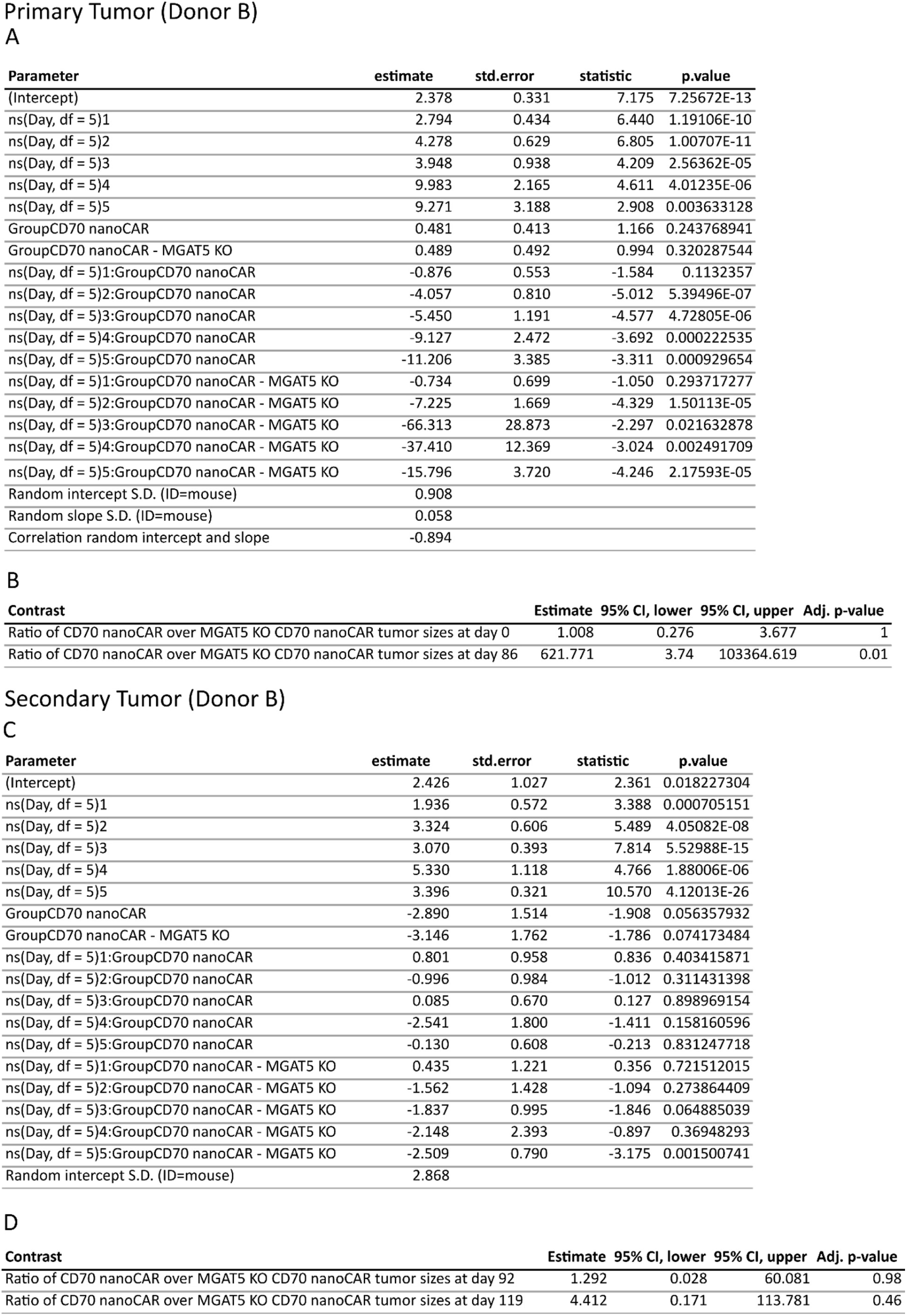
Longitudinal analysis of Donor B. Longitudinal analysis of the primary (A-B) and secondary tumor (C-D). A and C. A Tweedie GLMM (generalized linear mixed model) with log link was used to model the tumor volume. A. Summary of the model output, listing all parameter estimates for the model TumorVol ∼ ns(Day, df = 5)*Group + (Day | ID), ns = natural spline. The table gives parameters and standard errors on the natural log scale together with test statistics and multiple testing adjusted p-values (null hypothesis: parameter equal to zero). B and D. Inference for different research questions. In this table, the estimates and confidence intervals are transformed back to the original scale so we can interpret them in a straightforward way. Note: the estimate of 621.771 in panel B should be interpreted with caution, since within this contrast we are comparing to a tumor volume reaching zero. S.E.: Standard Error. CI: Confidence Interval.

**Supplementary Figure 6.**
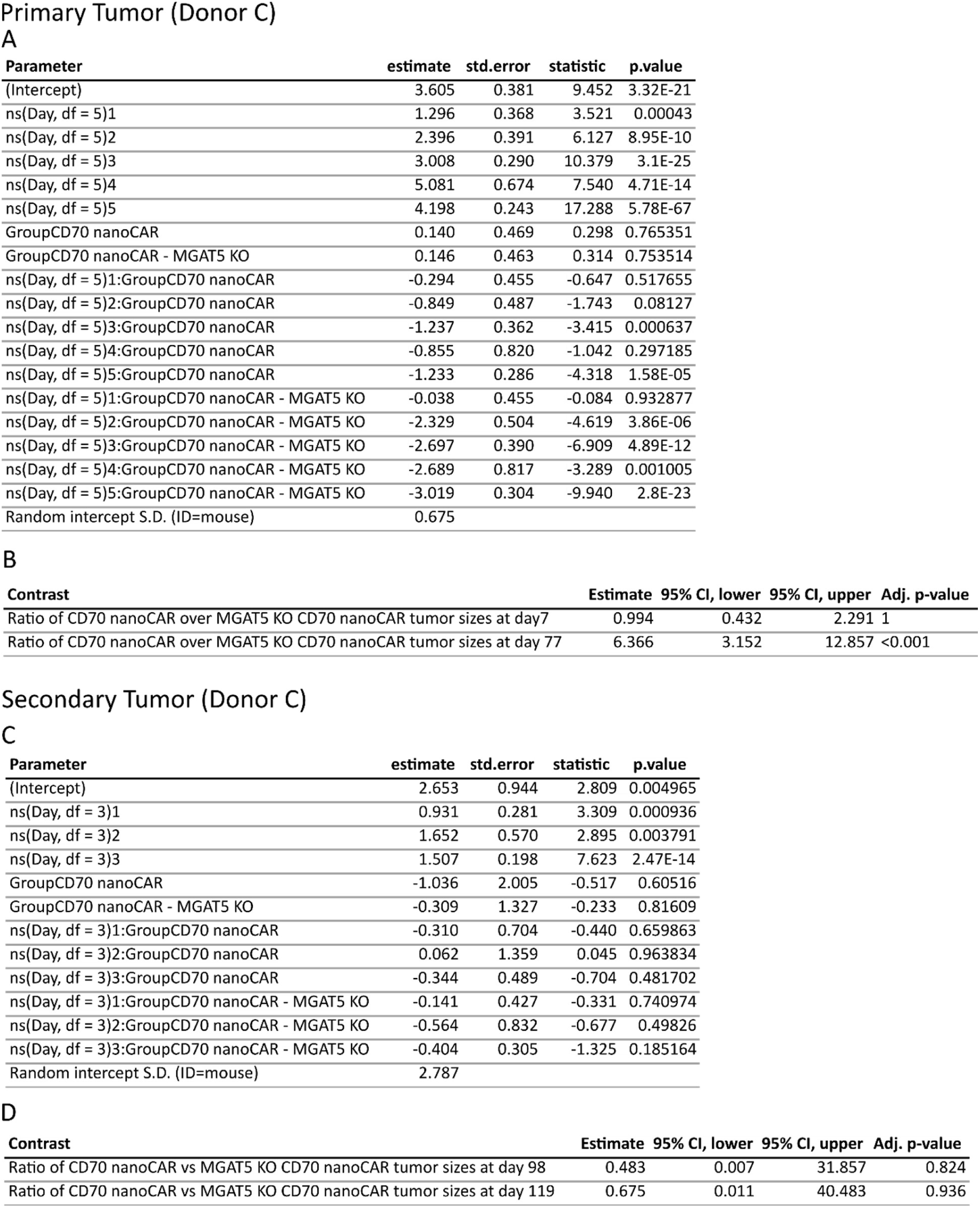
Longitudinal analysis of Donor C. Longitudinal analysis of the primary (A-B) and secondary tumor (C-D). A Tweedie GLMM (generalized linear mixed model) with log link was used to model the tumor volume. A and C. Summary of the model output, listing all parameter estimates for the model TumorVol ∼ ns(Day, df = 5)*Group + (Day | ID), ns = natural spline. The table gives parameters and standard errors on the natural log scale together with test statistics and multiple testing adjusted p-values (null hypothesis: parameter equal to zero). B and D. Inference for different research questions. In this table, the estimates and confidence intervals are transformed back to the original scale so we can interpret them in a straightforward way. S.E.: Standard Error. CI: Confidence Interval.

**Supplementary Figure 7.**
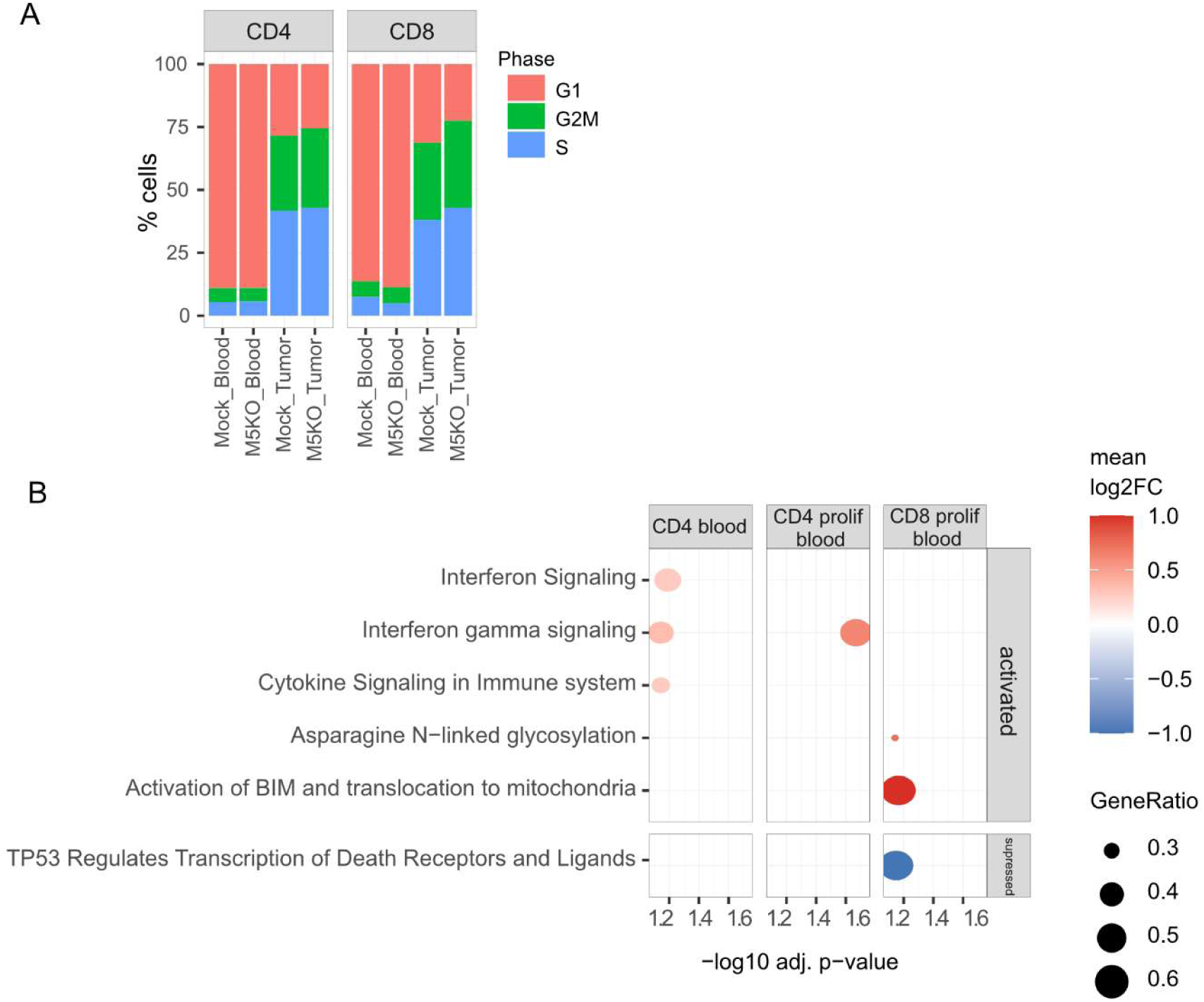
Single cell RNA sequencing of tumor-infiltrating and circulating CAR T cells. A. Both CD4^+^ and CD8^+^ subpopulations of MGAT5 KO CD70 nanoCAR T cells infiltrating the tumor showed a higher proportion of proliferating cells (G2M and S phase). B. Gene set enrichment analysis of Reactome pathways for CD4^+^ and CD8^+^ MGAT5 KO CD70 nanoCAR T cells circulating in blood.

## Supplementary Tables

Supplementary Table 1– List of differentially expressed genes in nanoCAR T cells from the 7 different clusters

Attached as separate file

Supplementary Table 2-Output of the Gene set enrichment analysis of Reactome pathways from the 7 different clusters

Attached as separate file

**Supplementary Table 3.** Overview of the different donors used in this study

| Donor | Experiment type in which T cells were used |  |
| --- | --- | --- |
| A | *in vivo* analysis tumor control efficacy SKOV-3 model |  |
| B | *in vivo* analysis tumor control efficacy SKOV-3 model |  |
| C | *in vivo* analysis tumor control efficacy SKOV-3 model |  |
| D | *in vivo* analysis tumor control efficacy Raji model |  |
| E | *in vivo* analysis tumor control efficacy Raji model |  |
| F | *in vivo* analysis tumor control efficacy Raji model |  |
| G | *in vivo* analysis tumor control efficacy Raji model | Cytokine-independent growth assay |
| H | analysis *ex vivo* tumor cell killing activity |  |
| I | analysis *ex vivo* tumor cell killing activity |  |
| J | scRNA seq experiment |  |
| K | degranulation assay |  |

**Supplementary Table 4.**
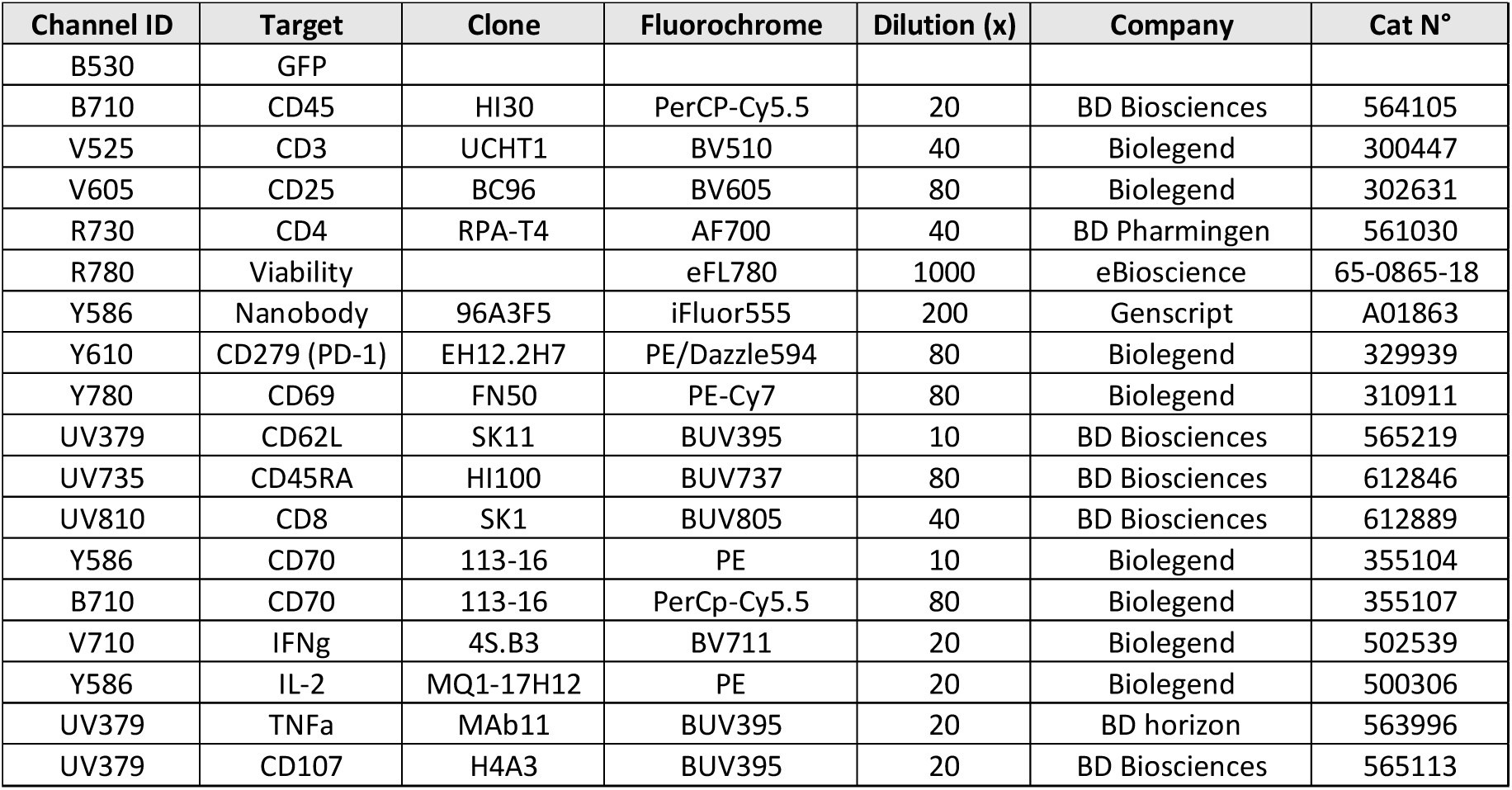
Overview of antibodies used in this study

| Channel ID | Target | Clone | Fluorochrome | Dilution (x) | Company | Cat N° |
| --- | --- | --- | --- | --- | --- | --- |
| B530 | GFP |  |  |  |  |  |
| B710 | CD45 | HI30 | PerCP-Cy5.5 | 20 | BD Biosciences | 564105 |
| V525 | CD3 | UCHT1 | BV510 | 40 | Biolegend | 300447 |
| V605 | CD25 | BC96 | BV605 | 80 | Biolegend | 302631 |
| R730 | CD4 | RPA-T4 | AF700 | 40 | BD Pharmingen | 561030 |
| R780 | Viability |  | eFL780 | 1000 | eBioscience | 65-0865-18 |
| Y586 | Nanobody | 96A3F5 | iFluor555 | 200 | Genscript | A01863 |
| Y610 | CD279 (PD-1) | EH12.2H7 | PE/Dazzle594 | 80 | Biolegend | 329939 |
| Y780 | CD69 | FN50 | PE-Cy7 | 80 | Biolegend | 310911 |
| UV379 | CD62L | SK11 | BUV395 | 10 | BD Biosciences | 565219 |
| UV735 | CD45RA | HI100 | BUV737 | 80 | BD Biosciences | 612846 |
| UV810 | CD8 | SK1 | BUV805 | 40 | BD Biosciences | 612889 |
| Y586 | CD70 | 113-16 | PE | 10 | Biolegend | 355104 |
| B710 | CD70 | 113-16 | PerCp-Cy5.5 | 80 | Biolegend | 355107 |
| V710 | IFNγ | 4S.B3 | BV711 | 20 | Biolegend | 502539 |
| Y586 | IL-2 | MQ1-17H12 | PE | 20 | Biolegend | 500306 |
| UV379 | TNFα | MAb11 | BUV395 | 20 | BD horizon | 563996 |
| UV379 | CD107 | H4A3 | BUV395 | 20 | BD Biosciences | 565113 |

